# Chemerin increases T-cell mediated cytotoxicity of human tumors via modulation of a novel CMKLR1/PTEN/PD-L1 axis

**DOI:** 10.1101/787697

**Authors:** Keith Rennier, Ethan Krug, Gurpal Virdi, Woo Jae Shin, Russell Pachynski

## Abstract

Chemerin (*RARRES2*) is an endogenous leukocyte chemoattractant known to recruit innate immune cells through its chemotactic receptor, CMKLR1. *RARRES2* is often downregulated in prostate cancer and across multiple tumor types compared to normal tissue. Additionally, a methylome-wide analysis identified *RARRES2* as one of the most hypermethylated genes in sarcoma. Recent studies have shown that augmenting chemerin in the tumor microenvironment significantly suppresses tumor growth, at least in part by recruitment of immune effector cells. However, as tumor cells have been shown to express functional chemokine receptors that can impact their phenotype, we hypothesized that chemerin might have additional, tumor-intrinsic effects. Here, we show for the first time that human cancer cells exposed to exogenous chemerin significantly upregulate PTEN expression and activity. Additionally, chemerin-induced PTEN expression correlated with a concomitant reduction in PD-L1 mRNA and protein expression. Chemerin treatment of tumor cells led to significantly reduced tumor cell migration/invasion, as well as significantly increased cytotoxicity by T cells. siRNA knockdown of tumor expressed CMKLR1 abrogated chemerin-induced modulation of PTEN and PD-L1 expression and activity, supporting the presence of a CMKLR1/PTEN/PD-L1 signaling cascade. We then compared chemerin treatment to PD-L1 inhibition by siRNA knockdown or the antagonistic antibody atezolizumab in T cell cytotoxicity assays and surprisingly found chemerin treatment was as effective as both PD-L1 knockdown and atezolizumab treatment in mediating tumor lysis. Collectively, our data show a novel link between chemerin, PTEN and PD-L1 in human tumor lines, with functional consequences that may have a role in improving T cell-mediated immunotherapies.

## Introduction

Successful immunotherapy requires activation of immune effector components to overcome mechanisms of tumor immune suppression and escape. Thus, targeting the various mechanisms by which tumors circumvent a successful immune response is critical for improved immunotherapy efficacy. In the local tumor microenvironment (TME), a skewed ratio of immunosuppressive leukocytes to their anti-tumor counterparts can lead to progressive tumor growth and resistance to immunotherapy ^1^ ^2^. Coordinated, selective recruitment of effector leukocytes - and optimized functional status - is thus central to peak immune response and immunosurveillance. Across several tumor types, preclinical models and human studies have shown that the number of tumor-infiltrating immune effector cells can positively correlate with survival ^3^, suggesting strategies that augment effector cell trafficking may improve immunotherapy response rates.

Chemerin, or *RARRES2* (retinoic acid receptor responder 2), is an endogenous leukocyte chemoattractant, but has myriad roles in adipogenesis, metabolism, angiogenesis, microbial defense, and cancer ^4–7^. Chemerin recruits innate immune cells along its concentration gradient to sites of inflammation via its G-protein coupled receptor (GPCR) chemokine-like receptor-1 (CMKLR1, aka ChemR23) ^8^ ^9^. Two additional GPCR receptors, CCRL2 and GPR1, for chemerin have been described in less detail, but have not been found to mediate significant signaling ^10, 11^. In humans, CMKLR1 expression on leukocytes has been shown in macrophages, dendritic cells (DCs), and NK cells with comparable expression in the mouse ^9, 12–14^. However, expression has also been seen in non-lymphoid tissues such as lung, skin, adipose tissue, vascular smooth muscle cells, and endothelial cells ^15^. While data is limited, CMKLR1 expression has been detected on human tumor cells ^16, 17^, suggesting that interaction with its endogenous ligand chemerin may modulate tumor cell phenotype, as seen with other chemokine/receptor pairs ^18^.

Chemerin/RARRES2 is commonly downregulated across several tumor types, including melanoma, prostate, and sarcoma, compared to their normal tissue counterparts ^6, 19–23^. Capitalizing on this, our group was the first to show that forcible re-expression of chemerin in the TME was able to recruit and increase tumor-infiltrating effector leukocytes, resulting in a significant reduction in the growth of aggressive B16 melanoma in a mouse model ^23^. While modulation of immune effector mechanisms is important, tumor cell-intrinsic oncogenic signaling pathways that control tumor cell growth can also significantly impact these directed immune responses, and thus play a key role in determining therapeutic efficacy.

PTEN (phosphatase and tensin homolog deleted on chromosome 10) is a dual-specificity phosphatase and critical tumor suppressor whose expression is downregulated and/or lost in many tumor types ^24^. PTEN loss has correlated with activation of the PI3K-AKT pathway, which is implicated in the pathogenesis of these cancers, and is particularly relevant in prostate cancer ^25–28^. Deleterious PTEN alterations are found in up to ∼20-30% of primary prostate cancer tissues and in ∼40-60% of metastatic tissues, and are among the most common genomic events in prostate cancer ^29–31^. While less commonly mutated in sarcoma, PTEN *downregulation* has also been shown to play an important role in a subset of soft tissue sarcomas (STSs), with one study showing 57% of STSs with decreased PTEN expression ^32^. Futhermore, aberrations in the downstream PI3K/Akt pathway are almost always implicated in the pathogenesis of sarcomas, with essentially 100% of advanced stage osteosarcomas showing dysregulation in this pathway ^33^.

Preclinical modeling, and now recent translational data show that PTEN loss – including known cell-intrinsic oncogenic effects – can lead to not only immune suppression within the TME, but resistance to immunotherapy as well ^34–36^. Thus, increasing PTEN and/or attenuating PI3K/Akt pathway activation may represent a promising strategy to improve overall immunotherapy success.

Here, we examine the effects of chemerin on tumor cell-intrinsic phenotype and describe – for the first time to our knowledge – the ability of chemerin to upregulate the expression and function of PTEN in human prostate and sarcoma tumor cell lines. Importantly, we show – also for the first time – that chemerin treatment of tumor cells results in a concomitant downregulation of surface PD-L1 expression, which directly translates into significantly improved T cell-mediated cytotoxicity, comparable to a currently approved PD-L1 checkpoint inhibitor, atezolizumab. These effects were dependent on CMKLR1, as siRNA knockdown completely abrogated these effects. Collectively, these studies suggest that subsequent to recruitment of effector leukocytes into the TME, chemerin may have the additional role of upregulating PTEN expression/function, thus suppressing PD-L1 expression, ultimately rendering tumor cells more susceptible to T cell mediated immunotherapies.

## Materials and Methods

### Cell Culture and Reagents

Cell lines were obtained from ATCC. DU145 (Human Prostate Cancer, HTB-81), PC3 (Human Prostate Cancer, CRL-1435) cells were cultured using RPMI 1640+10% Fetal Bovine Serum (FBS, Sigma)/1% L-glutamine (L-glut)/ 2% Penicillin/Streptomycin (P/S) at 37°C. SKES-1 (Human Ewing Sarcoma, HTB-86) and U2-OS (Human Osteosarcoma, HTB-96) cells were cultured with McCoy’s 5A+10% FBS/1% L-Glutamine/ 2% P/S. Exogenous recombinant human chemerin (2325-CM, R&D Systems) was added at varying concentrations (3nM or 6nM) for 48 hours.

### siRNA Transfection

Cells were plated at 100K cells/well overnight in 2mL of media. After 24h incubation, X-tremegene siRNA transfection reagent (#4476093001, Roche) and each siRNA at labeled concentrations were mixed and added drop-wise to the cell media. The optimal ratio of transfection reagent to siRNA (4:10) confirmed signal knockdown via Western blot. ChemR23/CMKLR1 (sc-44633, Santa Cruz), PD-L1 (sc-39699, Santa Cruz), and PTEN (6251S, Cell Signaling) siRNA were used as indicated in the figure legends.

### Flow Cytometry

∼300k cells/sample were blocked using 2uL mouse serum/200uL PBS for 10 minutes at 4°C. Each antibody (R&D Systems, 0.5 mg/mL) was added at 1uL/100k cells for 30 minutes at 4°C. Cells were washed with PBS + 1% FBS and centrifuged at 300g for 5 minutes. Cell were fixed with 10% buffered formalin and analyzed using a FACScalibur (BD Biosciences). Anti-human antibodies were as follows: hCMKLR1 PE (FAB362P, R&D), hPD-L1 FITC (clone MIH6, Invitrogen), hPD-1 A488 (clone EH12.2H7, Biolegend), hCTLA-4 PE (clone BNI3, Biolegend), hTIM-3 BV421 (clone F38.2E2, Biolegend), and IFN-γ APC (clone 4S.B3, Biolegend).

### Real-time RT-PCR

Sample RNA was isolated using Trizol (Invitrogen) and RNeasy Mini RNA Isolation kit (Qiagen). RNA concentrations were verified using NanoDrop 2000 (Thermo). RNA was converted into cDNA templates using iScript Advanced cDNA Synthesis kit (Bio-Rad). For real-time RT-PCR, cDNA templates were amplified with iTaq Universal SYBR Green Supermix (Bio-Rad) according to the manufacturer’s protocol. A CFX96 Real-Time PCR system (Bio-Rad) was used to quantify gene expression via the 2^ΔΔCt^ analysis method. Primer sequences were developed using Primer-Blast software (https://www.ncbi.nlm.nih.gov/tools/primer-blast/). Each RT-PCR result was corrected for GAPDH loading control using the same cDNA sample. Target qPCR primers for both 5’ - 3’ forward (Fwd) and reverse (Rev) sequences were as follows: Human PTEN (Fwd) – TGTTCAGTGGCGGAACTTGC, Human PTEN (Rev) – CCGTCGTGTGGGTCCTGAAT, Human PD-L1 (Fwd) – GCATTTGCTGAACGCCCCA, Human PD-L1 (Rev) – TGACTTCGGCCTTGGGGTAG, Human EGR-1 (Fwd) – TGAGCATGACCAACCCACCG, Human EGR-1 (Rev) – CTGCTGTCGTTGGATGGCAC, Human SRF (Fwd) – TCAGCAAGAGGAAGACGGGC, Human SRF (Rev) – CATGGGCTGCAGTTTTCGGG, Human GAPDH (Fwd) – GGCAAATTCCATGGCACCGT, Human GAPDH (Rev) – ACGTACTCAGCGCCAGCATC.

### Immunoblot Analysis

A RIPA buffer lysis solution plus protease and phosphatase inhbitior cocktail (Thermo) was used to lyse cells post experiment. Protein concentration was calculated using Pierce BCA Protein Assay kit (Thermo) based on manufacturer’s protocol. Bolt 4-12% Bis-Tris SDS-PAGE gels (Invitrogen) were loaded with equal sample protein amounts (50ug/sample). Gels were transferred to NitroBind Nitrocellulose membrane (Thermo). Rabbit anti-human PTEN (138G6), SRF(D71A9), EGR-1(44D5), pAKT(ser473, D9E), pS6(ser235/236, D57.2.2E), RPS6(5G10), PD-L1(E1L3N) was used to detect protein expression, w/ rabbit anti-human GAPDH HRP (D16H11, Cell Signaling) as loading control. Mouse anti-human AKT1 (094E10, Biolegend) was detected using anti-mouse HRP(poly4053, Biolegend). All antibodies were illuminated using Thermo SuperSignal West Dura per manufacturer’s protocol. Imaging was done using BioRad Molecular Imager ChemiDoc XRS+ System. Expression results were quantified using the BioRad Image Analysis Software.

### Immunoprecipitation and Phosphatase Activity

Cell lysis was collected as described above using IP cell lysis buffer (Thermo). Protein loading volumes were all normalized (200μg/sample). Anti-human PTEN (138G6, Cell Signaling, 1:250) was added to each protein sample and gently rotated overnight at 4°C. Next, PTEN-antibody solution were mixed with protein A/G agarose beads for 4 hours at room temperature. PTEN protein was released from beads using IgG elution buffer to examine phosphatase activity. To initiate, 3pM PIP_3_ (Echelon Biosciences, DiC8) was added to each PTEN-IP protein sample (200μg/sample) for 2 hours at 37°C. To measure free phosphate, the Malachite Green Phosphate Detection kit (12776, Cell Signaling) was followed according to manufacturer’s protocol. Absorbance values at 620nm direclty correlated to the amount of phosphatase activity in each sample.

### Tumor migration/invasion assay

A 24-well plate transwell inserts (6.5 mm, Costar, 8 μm pores) were pre-coated with 35 μl of 1 mg/mL matrigel (BD Biosciences) at 37°C for 2h. 0.5 × 10^5^ cells of each sample in serum-free medium were plated in the upper chamber and media with 10% FBS was added to the bottom well. After 24h, the non-invaded cells still in matrigel were removed by cotton swabs. The inserts were fixed in 100% methanol and stained with 0.1% crystal violet. Migrated cells were lysed with 10% acetic acid. Each absorbance was measured correlating to the amount of migrated cells per transwell insert (Bio-Tek Instruments).

### T cell-mediated cytotoxicity

Human T cells were isolated from donor PBMCs using Mojosort Human CD3 Isolation kit (Cat. #480022, Biolegend) according to manufacturer’s protocol. T cells were left untreated (naïve T cells), treated with IL-2 (100U/mL, R&D) or IL-2 + ImmunoCult CD3/CD28/CD2 T cell activation tetramers (activated T cells, 25uL/mL, #10970, StemCell). For cytotoxicity studies, DU145 cells were stained with CFSE (1μL/mL, #423801, Biolegend) according to manufacturer’s recommendation. CFSE positive DU145s were incubated with naïve (untouched) or activated T cells overnight (1:3 E:T ratio). Cells were stained with 7-AAD (5uL/1e6 cells, #420404, Biolegend) to identify dead cells via flow cytometry. Percent lysed was the fraction of DU145 cells that stained positive for both CFSE and 7-AAD. Ran triplicates for each sample set per experiment, n=4.

### RNA In Situ Hybridization and Image Analysis

Manual chromogenic RNAScope was performed with RNAScope 2.5 HD Reagent kit – brown from Advanced Cell Diagnostics (ACD, #322310), using optimized company protocols to detect target RNA at single cell level. Single ISH detection for PTEN (ACD Probe: 408511), PD-L1 (CD274 – ACD Probe: 600861), Positive Control Probe (PPIB - ACD Probe: 313901) and Negative Control Probe (Dapb - ACD Probe: 310043) was performed according to manufacturers’ protcols, Quantitative analysis was completed using regions of interest (ROIs) and by random sampling. Three comparable ROIs for each respective sample set were analyzed using HALO Software (3 ROIs per sample, repeated for n = 3 independent experiments).

### Statistics

All experiments were done independently, at least three times. Each time replicates of samples were prepared, collected, and analyzed independently. Means and standard errors of the mean (SEM) were calculated. Paired Student t-tests were used for comparison between two groups in each experiment. One-way ANOVA was used to compare more than two groups, including a post-hoc Tukey test to confirm where the differences occurred between groups. A p-value of less than 0.05 was considered statistically significant calculated using Microsoft Excel and GraphPad Prism v.8 software.

## Results

### Chemerin exposure can induce PTEN expression in tumor cell lines

We initially questioned whether chemerin, given its myriad roles, would have an impact on tumor intrinsic cell functions. Our previous studies in mouse models did not show detectable levels of CMKLR1 on mouse tumor cell lines, nor direct effect of recombinant chemerin exposure on tumor cell phenotype measured (eg proliferation, surfacer marker, etc) ^23^ . Given the prominent role of PTEN dysregulation in prostate and sarcoma tumors, we decided to initially study these tumor types using human cell lines. We looked at human tumor lines that had detectable CMKLR1 protein expression and genetically intact PTEN (DU145, U2OS, SKES) and used a CMKLR1+, PTEN null (-/-) cell line (PC3) as a control. Both prostate and sarcoma cell lines were analyzed for expression of CMKLR1, and showed detectable levels of CMKLR1 protein at both the intracellular and cell surface levels (supplementary figure 1). Cell lines had no detectable chemerin expression (anti-human chemerin ELISA assays, data not shown). We then investigated the effect of exogenous, recombinant chemerin on these cell lines. Chemerin is found systemically in plasma and engages CMKLR1 at low nanomolar concentrations ^8^, thus we chose to initially focus in this range of concentrations. Cell lines were were plated as indicated and incubated with complete media containing 6 nM recombinant chemerin protein or PBS (the diluent control) for 48 hours. Culture media with PBS or chemerin was changed daily. Following treatment, PTEN mRNA expression was then quantified using quantitative real-time RT-PCR analysis. PTEN expression was normalized to GAPDH as a housekeeping gene for each sample. Here, we found PTEN mRNA expression was significantly upregulated over control-treated cells in all PTEN WT cell lines tested (figure 1A, B). As expected, there was no detection of PTEN in the PC3 cells, while we saw an approximately 2-fold increase in PTEN expression in the DU145 cells, and a ∼2.5-fold increase in the sarcoma SKES and U2OS cells after incubation with chemerin.

**Figure 1.**
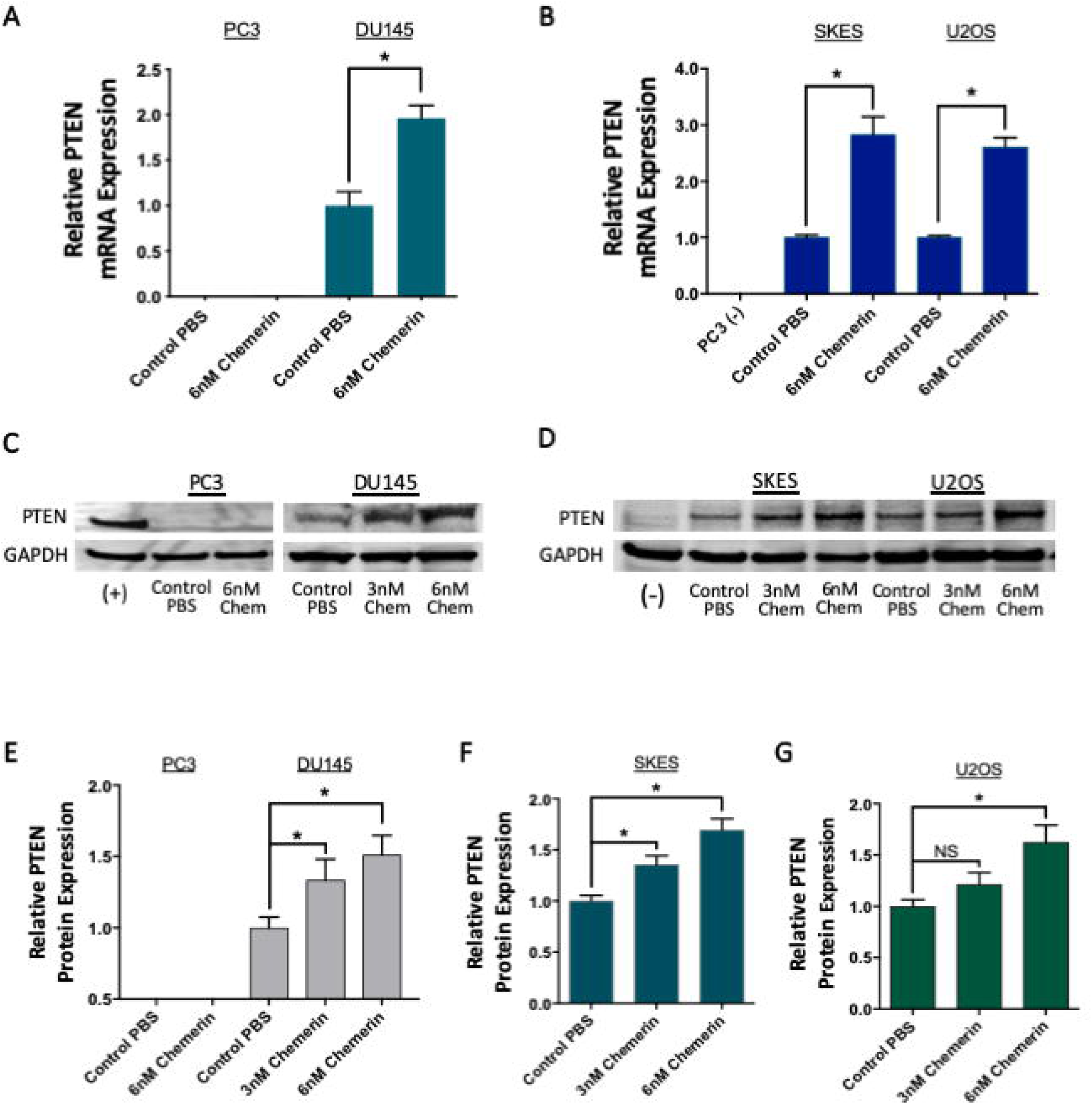
**Recombinant chemerin upregulates PTEN expression in tumor cells.** A. Real time RT-PCR results of PTEN mRNA expression in prostate cancer cells treated with vehicle control (PBS) or 6nM recombinant chemerin (6nM Chem). PTEN Expression is normalized to GAPDH loading control for each sample and normalized to Control PBS across the dataset(\**P* < 0 01, n = 4). B. Real-time RT-PCR results of PTEN mRNA expression in Ewing sarcroma (SKES) and osteosarcoma(U2OS) cells treated with PBS(Control) or 6nM recombinant chemerin. PTEN Expression is normalized to GAPDH loading control for each sampe and normalized to Control PBS across the dataset(\**P*< 0.01, n = 4). C. Representative Western blots for PTEN protein expression in Normal Prostate - RWPE1(+), PC3, and DU145 cells treated with vehicle (Control PBS) or 6nM Chem(6nM chemerin) for 48h. D. Representative Western blot for PTEN protein expression In PC3 (-), SKES, and U2OS cells treated with vehicle (Control PBS) or 3nM or 6nM Chem (6nM chemerin) for 48h. E. Quantified Western blot results showing PTEN protein expression in control or chemerin treated PCa cells. ormalized to GAPDH loading control for each respective sample and each dataset is normalize to Control PBS (\**P* < 0 05, n = 3) F-G. Quantified Western blot results for PTEN protein expression in PBS(Control) or Chemerin treated sarcoma cells (F. SKES, G. U2OS). Each sample is normalized to GAPDH loading control and the dataset is normalized to Control PBS. Each sample set was independently repeated three times(\**P*< 0.05, n = 3).

Next, we wanted to confirm these findings at the protein level, as mRNA expression does not perfectly correlate with protein production ^37^. To begin to investigate a dose-response relationship, we incubated cell lines with both 3 and 6nM recombinant chemerin and compared to the control group. Western blot analyses showed a noticeable upregulation of PTEN protein expression in all three cell lines after a 48h incubation, again with no detectable PTEN found in the PC3 cells (figure 1C, D). Quantification showed that PTEN expression increased with an increasing concentration of chemerin, suggesting a dose-response. PTEN protein was increased ∼1.5-fold in DU145 cells, and ∼1.6-1.7-fold in U2OS and SKES cells compared to the controls (figure 1E-G). Because PTEN has been shown to affect cell proliferation and apoptosis in a number of settings ^38–41^, we assessed the effects of exogenous chemerin on these parameters in our cells. Neither *in vitro* cell proliferation nor apoptosis (supplemental figure 2 and 3) was significantly altered after chemerin exposure over a 72 hour perioid, compared to controls. Collectively, these results show that exogenous chemerin significantly induces PTEN mRNA and protein expression in a dose-dependent manner, without significant impact on their *in vitro* proliferation or apoptosis.

### CMKLR1 mediates chemerin-induced PTEN expression and function

Given the other described chemerin receptors (CCRL2 and GPR1) are non-signaling ^10, 11^, we focused on the role of CMKLR1 in mediating chemerin-induced PTEN upregulation in these cell lines. Control or CMKLR1-specific siRNA were utilized in transient transfection experiments optimized with the siRNA transfection reagent. Knockdown of CMKLR1 protein was confirmed at both the intracellular and cell surface levels (supplemental figure 4). Compared to controls, significant increases in PTEN expression, by both qPCR and Western blot, were seen with exposure to 6nM chemerin during mock and control siRNA transfections in all cell lines. However, only with CMKLR1 siRNA was there a complete abrogation of chemerin-induced PTEN expression (figure 2A). This establishes the role of CMKLR1 in mediating the chemerin-induced increase of PTEN expression, at both the mRNA and protein levels.

**Figure 2.**
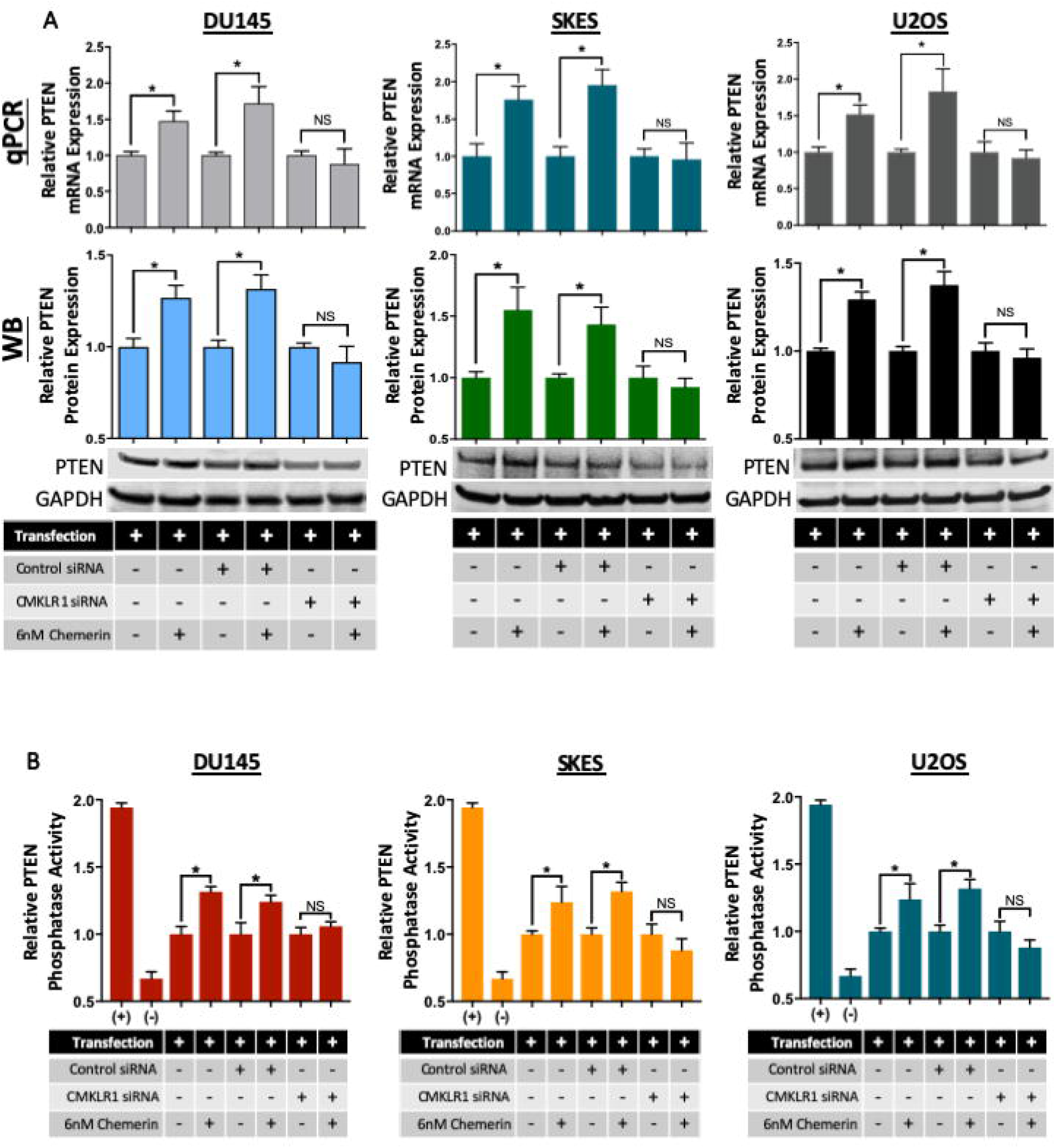
**Chemerin induces PTEN expression and activity via CMKLR1**. **A** (Top) Real-time RT-PCR results of PTEN mRNA expression in DU145(Left), SKES (Middle), U2OS (Right) cancer cells transfected with the following groups: Mock (no siRNA), Control siRNA (non-specific sequence), or CMKLR1 siRNA. Following transfection, each respective group was treated with PBS (Control) or 6nM recombinant chemerin. PTEN expression is normalized to GAPDH loading control for each sample and each pair was normalized to Control PBS, respectively (\**P* < 0.01, n = 4. NS = No significant difference). (Middle) Representative Western blot for PTEN protein expression in the transfected DU145, SKES, U2OS cell subsets treated with PBS or 6nM chemerin for 48h. (Bottom) Quantified Western blot results showing PTEN protein expression in PBS or Chemerin treated DU145, SKES, U2OS cells following transfectlon. Sample expression was normalized to GAPDH loading control and each pair was normalized to each Control PBS, respective y (\**P*< 0.05, n = 3). **B.** Cells were treated with either PBS or 6nM chemerin for 48h PBS or Chemerin DU145 cells transfected with Mock (no siRNA), Control siRNA, or CMKLR1 siRNA. Protein samples were collected after each experiment using an IP lysis buffer. PTEN was immunoprecipitated from 75 µg of protein in each sample to quantify specific phosphatase activity for each phosphatase assay, each sample was incubated with 3pM PIP_3_ in duplicate wells for 2h at 37°C. Next, wells were incubated with Malachite Green, and the absorbance was read, correlating to total amount of free phosphate (pMol) in solution/well. Each sample set and condition were repeated in three independent experiments(n = 3). Positive control (+) corresponds to 3pM PIP^3^ + recombinant human PTEN protein and negative control (-) was incubation with 3pM PIP^3^ only. Results show that 6nM chemerin increases PTEN phosphatase activity in Mock and Control siRNA transfected cells after 48h treatment. Although, this increase is lost following CMKLR1 knockdown via siRNA transfection. \**P* < 0.05, compared to each vehicle control (PBS) treated cells for each cell line and transfection subset. DU145 (Left), SKES (Middle), U2OS (Right).

While PTEN increased due to chemerin in all cell lines, we next wanted to determine if this PTEN increase was indeed functional. PTEN phosphatase activity modulates PI3K-induced phosphatidylinositol-3,4,5-triphosphate (PIP_3_), which is a critical factor in mediating subsequent signaling pathways involved in cell survival, proliferation, and migration ^42^. We next studied the ability of PTEN protein to dephosphorylate a key PI3K pathway constituent, PIP_3_ phosphate, following PBS or chemerin incubation. Protein lysates were collected following each 48h experiment for PTEN immunoprecipitation. PTEN phosphatase activity was significantly increased after 48hr chemerin exposure compared to control-treated cells (figure 2B), suggesting the chemerin-induced PTEN retained its ability to function as a phosphatase. Likewise, specific CMKLR1 knockdown – but not mock transfection nor control siRNA – completely abrogated the chemerin-mediated increase in PTEN phosphatase activity (figure 2B). Together, these findings confirm chemerin’s ability to induce significant *functional* PTEN expression via CMKLR1 in human tumor lines.

### Chemerin treatment, mediated by CMKLR1, significantly reduces tumor migration and invasion

PTEN has multiple roles in tumor suppression, including inhibition of tumor cell proliferation, invasion and migration ^43, 44^. While chemerin did not impact cell proliferation, we then decided to look further at other aspects of tumor cell phenotype. Using a tumor migration model, we examined the effects of chemerin treatment for 48h, in line with our previous experimental conditions. Control or chemerin-treated cells were allowed to migrate on matrigel for 24h, after which cells were fixed and stained for analysis. Imaging showed a noticeable decrease in tumor migration/invasion in all three of the cell lines treated with chemerin compared to control-treated cells (figure 3A). Quantification of migrated tumor cells shows that chemerin treatment significantly reduced tumor cell invasion by 29% in DU145 cells, 31% in U2OS cells, and 22% in SKES cells compared to control-treated samples, respectively (figure 3B).

**Figure 3.**
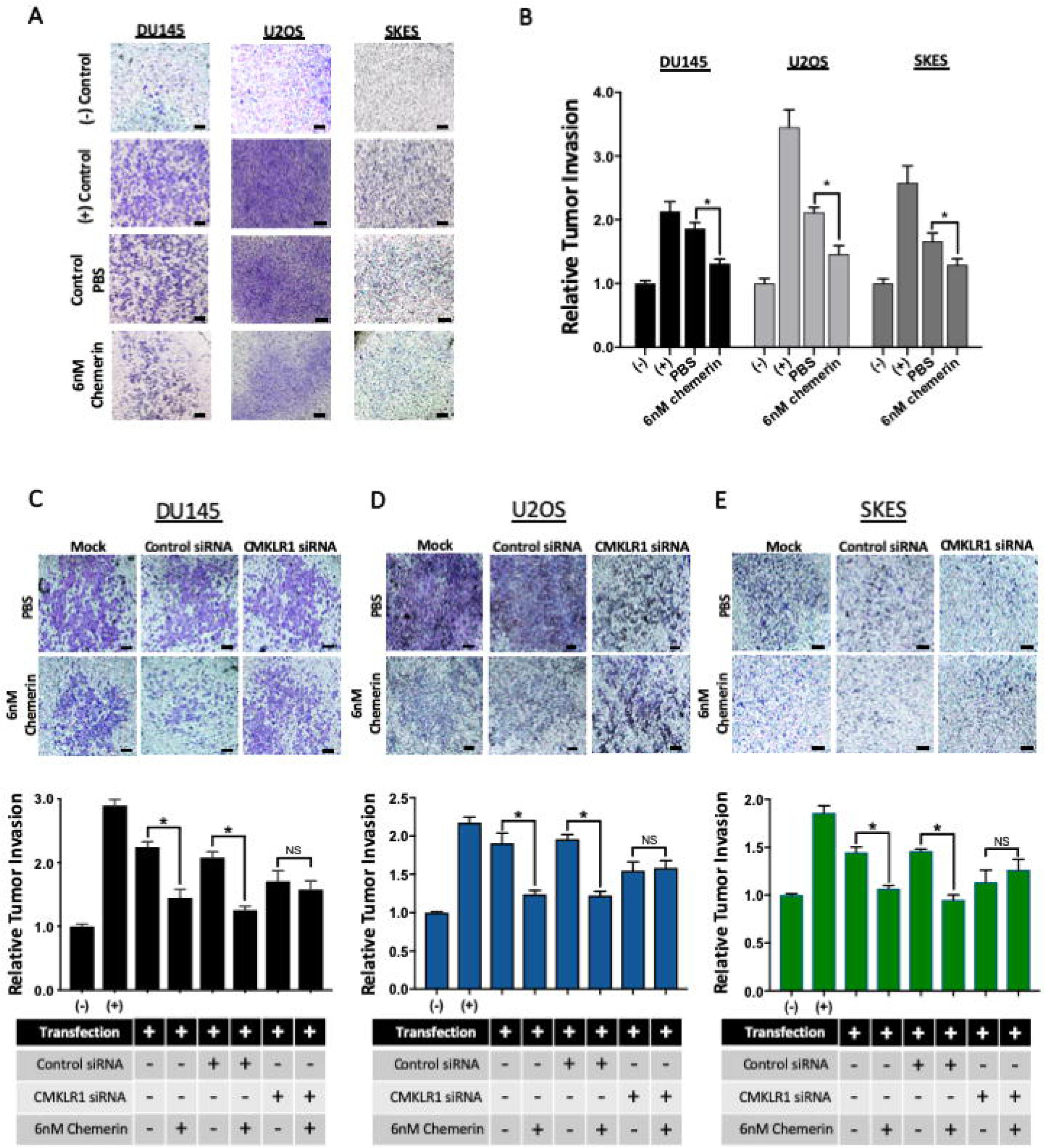
Chemerin treatment significantly reduces tumor cell migration and invasion in a CMKLR1-dependent manner. **A.** Representative 4X images showing tumor cell invasion normalized to baseline cell migration, No matrigel matrixand No FBS (n = 4). The following groups were compared: No matrigel - No FBS, No matrigel - CM + 10% FBS, 1mg/mL matrigel + cells treated with 48h PSS, or 1mg/mL matrigel + cells treated with 48h 6nM recombinant human chemerin (6nM chemerin). Scale bar = 100µm. **B.** Quantified tumor cell invasion results for each respective tumor cell line comparing matrigel invasion in cells treated with PBS (vehicle) or 6nM Chemerin for 48h. Following treatment, cells were allowed to migrate for 24h. Cells were fixed with methanol and stained with 0.1% Crystal violet. After Imaging, cells were lysed with 10% acetic acid. To quantify, each sample in duplicate wells absorbance was read at 590nm. Each respective absorbance is normalized to the baseline migration when using a no matrigel and no FBS control setup (\**p*< 0.05, n = 4). CMKLR1 knockdown negates chemerin’s ability to decrease DU145 cell invasion. **C-E (Top)**. Representative 4X Images showing tumor cell invasion of PBS or Recombinant Chemerin treated C) DU 145, D) U2OS, and E) SKES cells transfected with Mock(no siRNA), Control siRNA, or CMKLR1 siRNA. Each sample set and condition were performed in triplicate wells. Overall migration was normalized to baseline cell migration using No matrigel matrix and No FBS gradient(n = 3). Each of the following groups were compared: No matrigel - No FBS, No matrigel - CM + 10% FBS, 1mg/mL matrigel +cells treated with 48h PBS, or 1mg/mL matrigel + cells treated with 48h 6nM recombinant human chemerin (rHchemerin). Scale bar = l00µm. **F-H (Bottom).** Quantified tumor cell invasion results for each subset of transfected C) DU145, D) U2OS, and E) SKES cells. Each set was treated with either PBS (vehicle) or 6nM Chemerin for 48 h. Cells were processed as previously described, ran in duplicate wells and the relative sample absorbance was read at 590nm. Each respective sample absorbance is normalized to baseline migration using the no matrigel and no FBS control (\**P* < 0.05, n = 4. NS = No significant difference).

We next wanted to confirm that the effect of chemerin treatment on reducing tumor migration/invasion was mediated by CMKLR1, and not “off-target” effects. CMKLR1 siRNA knockdown experiments were performed as described above, using the matrigel invasion assay. Mock transfection and control siRNA cells continued to display significantly reduced tumor cell invasion after chemerin treatment, compared to control-treated tumor cells. In general, CMKLR1 siRNA knockdown decreased overall cell invasion in both control and chemerin-treated groups compared to mock transfection or control siRNA groups. However, CMKLR1 knockdown completely abolished the ability of chemerin to inhibit tumor cell migration and invasion comared to control-treated cells in all three lines (figure 3C-E). Collectively, these studies show a significant functional impact of chemerin treatment on tumor cell lines, and may represent a way to reduce tumor metastatic potential *in vivo*.

### Chemerin upregulates PTEN and concomitantly decreases PD-L1 expression on tumor cells, via CMKLR1

Recent evidence correlates PTEN expression and function to programmed death ligand-1 (PD-L1) expression in cancer, and this has been shown to be dependent on the PI3K pathway and S6 kinase (S6K) activation ^45–48^. Thus, we decided to look at PD-L1 expression in the context of chemerin exposure and PTEN expression. We tested a wide range (3-62nM) of recombinant chemerin on DU145 tumor cells, and then assessed for both PTEN and PD-L1 expression. As previously seen, 6nM chemerin treatment produced the most robust increase in PTEN expression, with an obvious dose-response relationship seen. Importantly, we also saw a concomitant, significant decrease in PD-L1 mRNA expression via qPCR (figure 4A).

**Figure 4.**
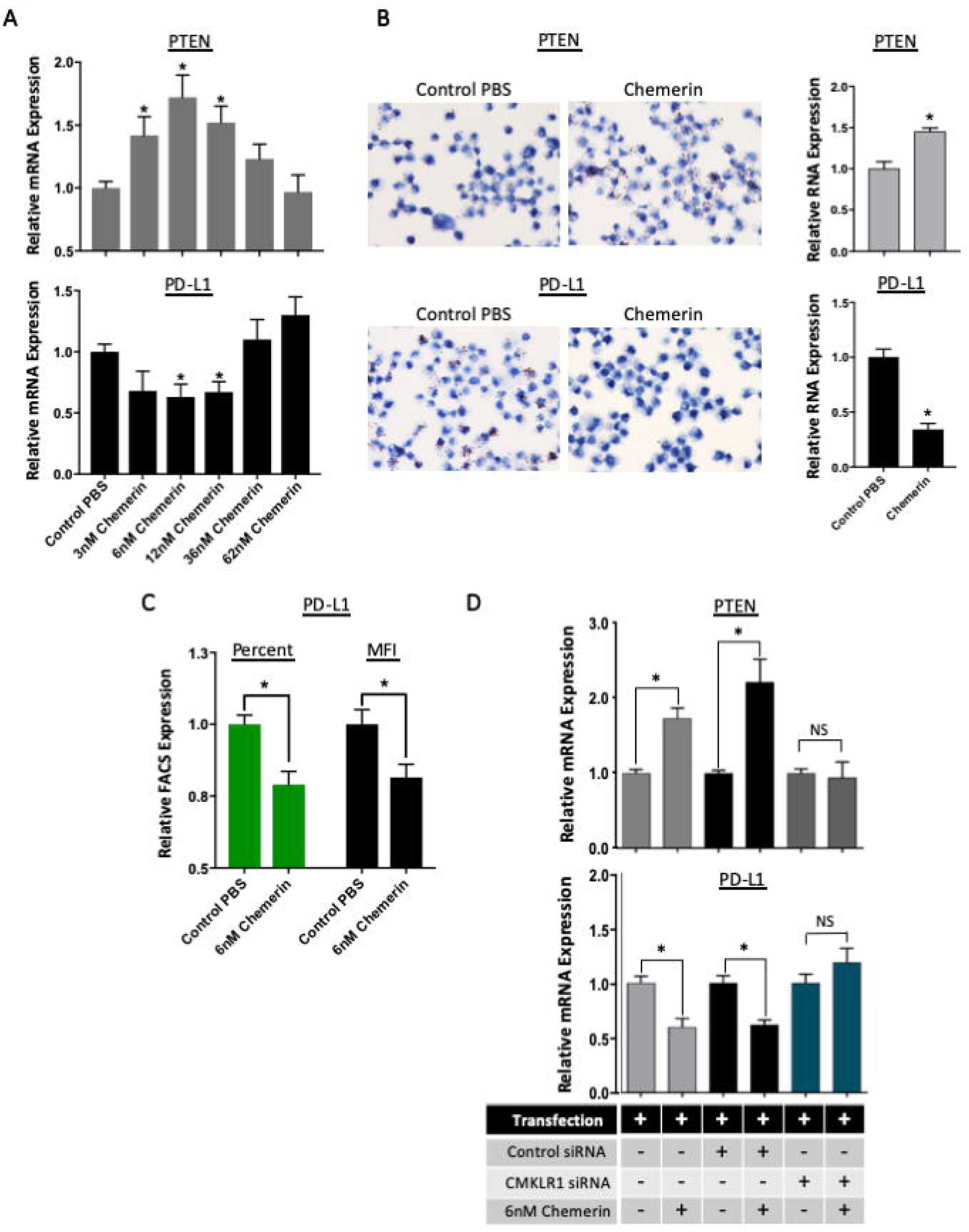
Chemerin upregulates PTEN with simultaneous decrease in PD-L1 via CMKLR1. Recombinant chemerin upregulates PTEN expression in tumor cells. **A.** Real-time RT-PCR results of PTEN mRNA expression in prostate cancer cells treated with vehicle control (PBS) or 6nM recombinant chemerin (6nM Chem). PTEN Expression is normalized to GAPDH loading control for each sample and normalized to Control PBS across the dataset (\**P* < 0.01, n = 4). **B.** Real-time RT-PCR results of PTEN mRNA expression in Ewing sarcroma (SKES) and osteosarcoma (U2OS) cells treated with PBS (Control) or 6nM recombinant chemerin. PTEN Expression is normalized to GAPDH loading control for each sample and normalized to Control PBS across the dataset (\**P* < 0.01, n = 4). **C.** Representative Western blots for PTEN protein expression in Normal Prostate - RWPE1 (+), PC3, and DU145 cells treated with vehicle (Control PBS) or 6nM Chem(6nM chemerin) for 48h. **D.** Representative Western blot for PTEN protein expression in PC3 (-), SKES, and U2OS cells treated with vehicle (Control PBS) or 3nM or 6nM Chem (6nM chemerin) for 48h. **E.** Quantified Western blot results showing PTEN protein expression in control or chemerin treated PCa cells normalized to GAPDH loading control for each respective sample and each dataset is normalize to Control PBS (\**P* < 0.05, n = 3). **F-G.** Quantified Western blot results for PTEN protein expression in PBS(Control) or Chemerin treated sarcoma cells (F. SKES, G. U2OS). Each sample is normalized to GAPDH loading control and the dataset is normalized to Control PBS. Each sample set was independent repeated three times (\**P* < 005, n = 3).

To confirm, we looked at RNA *in situ* hybridization (ISH) staining for PTEN and PD-L1. Human PTEN and PD-L1 RNA probes were used to quantify the total RNA expression in chemerin and control-treated cells. Image analysis software found that chemerin significantly upregulated PTEN and simultaneously decreased PD-L1 RNA expression (figure 4B). Additionally, we evaluated the impact of chemerin on tumor cell surface expressed PD-L1 protein, as this ultimately mediates its immunosuppressive effects. FACS staining analyses for PD-L1 revealed a signficiant *decrease* in surface expression in the chemerin-treated tumor cells compared to controls (figure 4D). We then wanted to investigate the role of CMKLR1 in the chemerin-mediated suppression of PD-L1 expression. We found that only knockdown of CMKLR1 – and neither mock transfection nor control siRNA – completed abrogated both the significant increases in PTEN and decreases in PD-L1 expression seen with chemerin treatment (figure 4E).

These data confirm a key inverse relationship between the tumor suppressor, PTEN, and a key immune checkpoint inhibitor, PD-L1, as previously described ^45–48^. More importantly, our findings show - for the first time - that chemerin can directly modulate this established PTEN/PD-L1 axis via CMKLR1 in human tumor cells.

### Chemerin activates transcription factors SRF and EGR-1 in correlation with PTEN upregulation

To further elucidate this novel pathway, we begin to investigate the underlying mechanisms of the chemerin-PTEN interaction. Recent research shows chemerin binding to CMKLR1 leads to transcriptional activation of the serum response factor (SRF) ^49^. Further, increased SRF expression induces activation of its target gene EGR-1, early growth response – 1, which directly regulates PTEN expression ^50^. Abberant PI3K pathway activation leads to a decrease in SRF levels and results in reduced binding to the EGR-1 promoter necessary for EGR-1 transcription ^51^. Thus, we hypothesized this signaling interplay may describe the pathway between chemerin and PTEN via CMKLR1 in DU145 cells. Therefore, we examined both SRF and EGR-1 expression in chemerin-treated DU145 cells as previously described above. In correlation with upregulated PTEN expression, our RT-qPCR results show a significant increase for both SRF (1.75-fold) and EGR-1 (1.91-fold) mRNA expression in the chemerin-treated DU145 cells compared to PBS alone (figure 5A). Similarly, western blot analysis show an substantial increase in SRF and EGR-1 protein expression, 1.68-fold and 1.57-fold increase respectively, directly correlating with increased PTEN protein expression (1.67-fold increase) (figure 5B, 5E-5G). Taken together, these results suggest chemerin binding CMKLR1 induces increased SRF and EGR-1 expression upstream of augmented PTEN expression and activity in DU145 cells.

**Figure 5.**
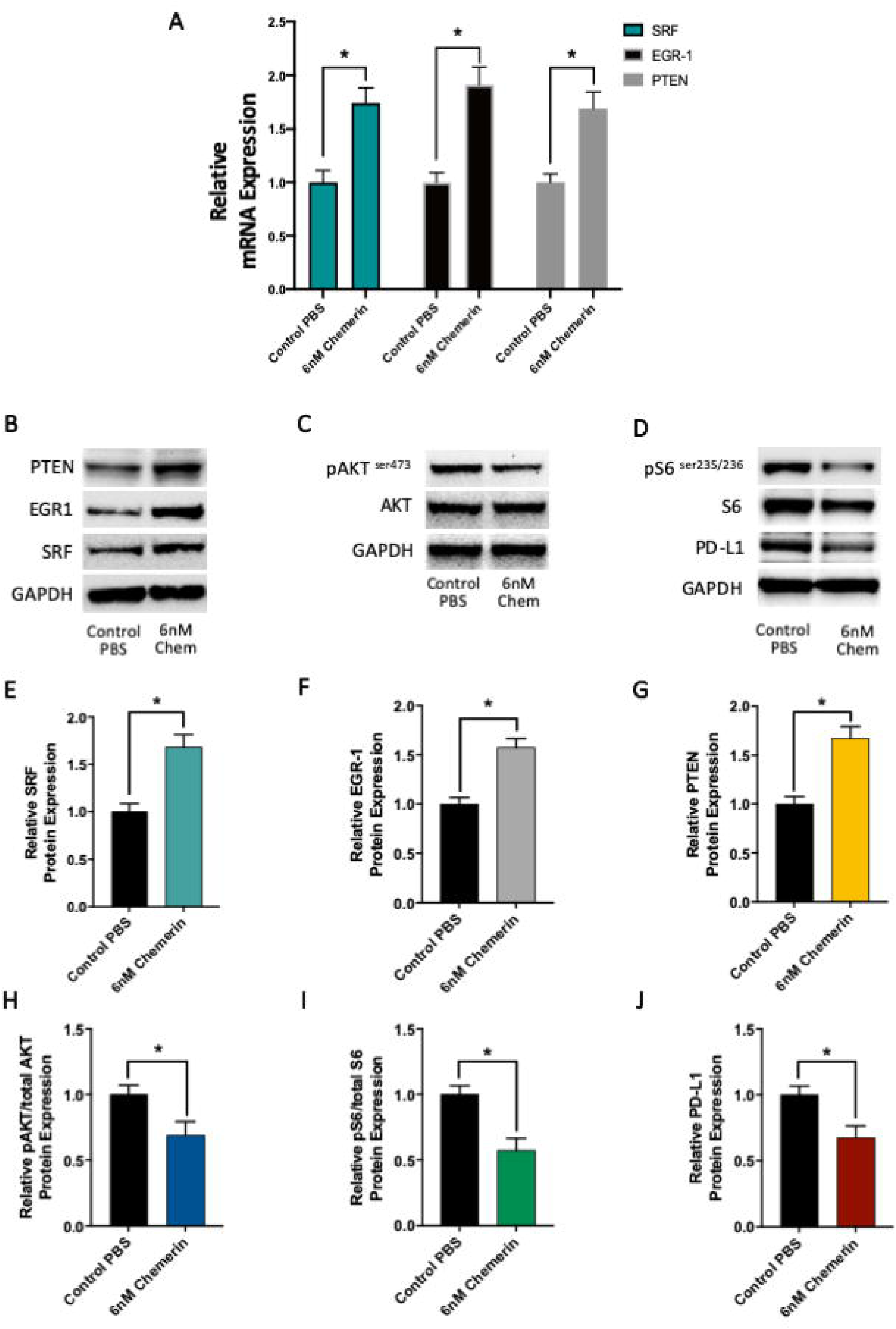
Chemerin modulates PTEN/AKT/PD-L1 and its signaling constituents. **A.** Quantitative real-time RT-PCR results of transcriptional regulators, SRF and EGR-1, including PTEN mRNA expression in DU145 cells treated for 48hrs using either vehicle control (Control PBS) or 6nM chemerin (\**P* < 0.05, n = 3). **B-D.** Representative Western blots for B) PTEN, SRF, and EGR-1 C) pAKT (ser 473), total AKT D) pS6, total S6, and PD-L1, each set including GAPDH loading control for DU 145 cells treated with vehicle (control PBS) or 6nM Chem (6nM chemerin) for 48h. **C. E-G.** Quantified Western blot results showing E) SRF F) EGR-1G) PTEN protein expression in control or chemerin treated DU145 cells. All samples were normalized to GAPDH loading control for each respective sample and each dataset is normalized to each respective Control PBS (\**P* < 0.05, n = 3). **H-J.** Quantified Western blot results for H) pAKT vs totalAKT J) pS6 vs Total S6 and J) PD-L1 protein expression in PBS (Control PBS) or 6nM chemerin treated DU145 cells. Each sample is normalized GAPDH loading control and the dataset is normalized to the respective Control PBS. Each sample set was independently repeated four times (\**P* < 0.05, n = 4).

### Chemerin inhibits pAKT and pS6 signaling leading to decreased PD-L1 expression

To further characterize the relationship between chemerin, PTEN and PD-L1, we examined the protein levels of p-Akt (ser473) and pS6 (ser235/236) by western blot. PTEN negatively regulates the PI3K/Akt pathway and overall PTEN activation inversely correlates with p-Akt expression ^25–28, 33^. Furthermore, PTEN loss or PI3k genetic alterations in prostate, breast, or glioma tumors show significantly augmented PD-L1 expression ^47, 52^. Lastwika et al. show that PD-L1 is tightly regulated by the Akt-mTOR pathway, where activation can lead to immune escape for some tumor types ^53^. Previous inhibition studies confirmed that the control of PD-L1 is PI3k/Akt/mTOR specific via mTORC1 (pAKT) and mTORC2 (pS6) signaling ^53, 54^. Thus, we studied the key PI3k/Akt/mTOR constituents in pAkt and RPS6, ribosomal protein S6, to confirm the downstream impact on PD-L1. Here, we again treat DU145s with either PBS or 6nM chemerin as previously described and examine protein expression via western blot. Figure 5C shows a noticeable p-Akt (ser473) decrease in protein expression following chemerin incubation, compared to the PBS treatment. Quantification of western blot data from 4 independent sample sets show a 29% decrease in pAkt (ser473) protein expression in chemerin-treated cells compared to the control PBS group (figure 5H). We also show that chemerin treatment leads to a substantial decrease in both pS6 (ser235/236) and PD-L1 protein expression compared to the control PBS group (figure 5D). Quantification of 4 independent sample sets reveal a 43% decrease in phospho-S6 and a 32% decrease in PD-L1 protein expression (figure 5I, 5J). Thus, our experimental results suggest that 48h chemerin exposure increases PTEN expression and activity leading to a subsequent negative regulation of the Akt – mTOR - PD-L1 signaling cascade. These results are consistent with previous studies of augmented PTEN expression on the PI3k/Akt/mTOR pathway and its signaling constituents ^25, 27, 28, 33, 35, 55^.

### Chemerin treatment significantly improves T cell-mediated cytotoxicity of tumor cells via CMKLR1

Our findings that chemerin can downregulate PD-L1 suggest it may play a role in T cell-mediated cytotoxicity. PD-L1 is known to inhibit T cell function via its interaction with programmed cell death-1 (PD-1) on T cells ^56^. To investigate, we isolated human T cells from donor PBMCs for cytotoxicity assays to target DU145 and U2OS tumor cells. Mirroring published data, we found that unstimulated, naïve T cells were only able to mediate low levels of DU145 cytotoxicity ^57, 58^. Both activation (with IL-2 and CD2/CD3/CD28 tetramers) and increasing effector to target (E:T) cell ratio resulted in improved tumor cell killing (supplemental figure 5). Importantly, we saw a significant increase in activated T cell-mediated cytotoxicity of DU145 cells that were treated with chemerin (48h), compared to controls (figure 6A). However, this effect was only seen at lower E:T ratios (i.e. 0.5:1 to 3:1), while the impact of chemerin treatment appeared to lessen at higher E:T ratios. This could suggest that larger numbers of activated T cells per tumor target cell act to obscure the effect mediated by chemerin treatment. Interestingly, prostate cancers typically are less infiltrated with immune cells compared to most other tumor types ^59^, suggesting that the lower E:T ratios in our assays are more physiologically relevant to the human TME. Furthermore, infiltrating T cells show an exhausted phenotype, as evidenced by high PD-1 expression and decreased IFN-γ ^60^ (supplemental figure 6), and thus PD-L1 expression on tumor cells is likely to modulate prostate-infiltrating T cell function.

**Figure 6.**
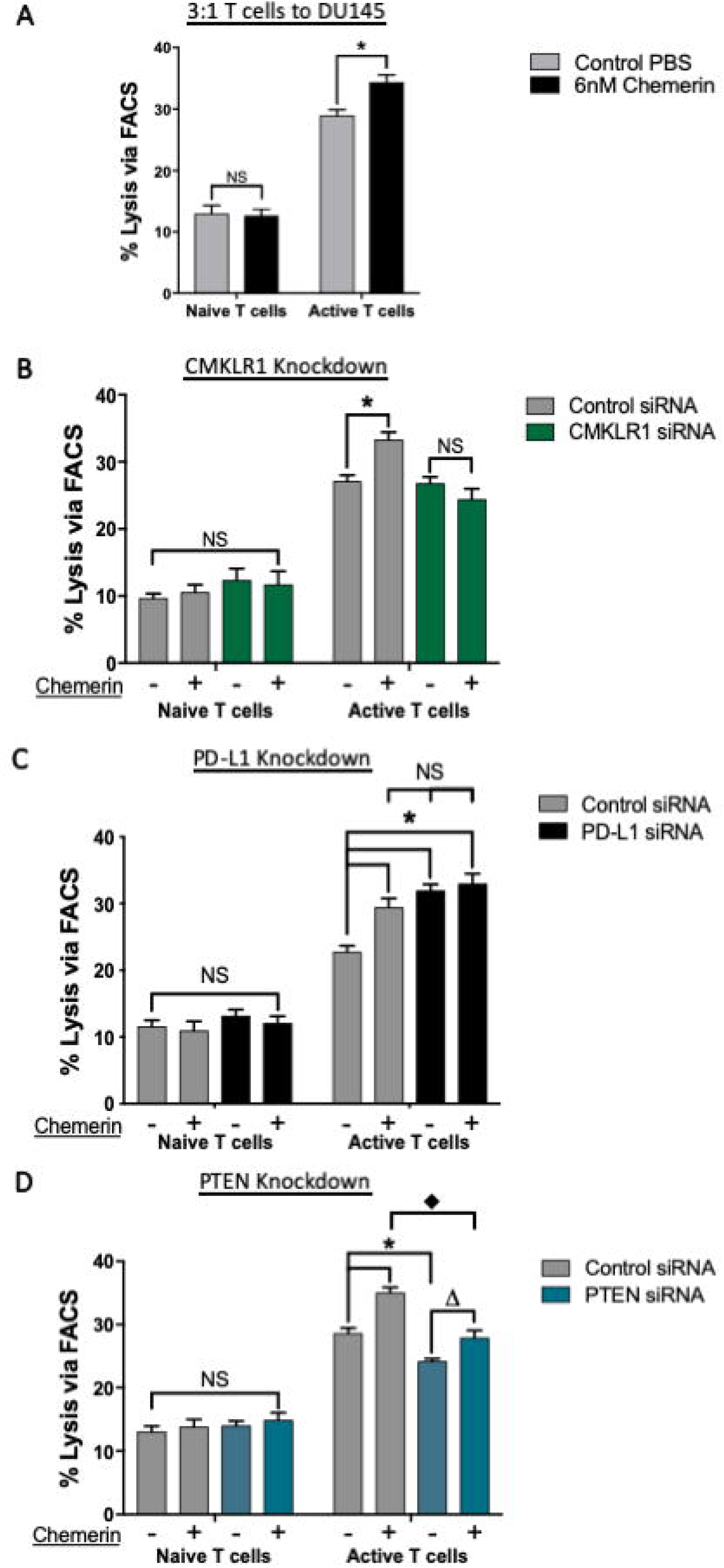
Chemerin improves T cell mediated cell cytotoxicity in tumor cells, and is mediated in part by PTEN and PD-L1. **A.** Naive and CD3-CD28-CD2 tetramer activated T cell mediated cytotoxicity in PBS vs 6nM chemerin treated DU 145s using the most effective ratio, 3:1 E:T(\**P* < 0.01, using triplicate samples for each experiment and repeated for n = 3 independent experiments). Chemerin treated DU145 cells resulted in a 6-8 percent increase in active T cell killing compared to PBS treated cells. **B.** T cell mediated cytotoxiclty in control vs CMKLR1 siRNA transfected DU145 cells. The effect of 6nM chemerin on cytotoxicity is abrogated following CMKLR1 knockdown (\**P* < 0.05, n = 3). **C.** Cytotoxicity in control vs PD-L1 siRNA transfected DU 145 cells. PD-L1 knockdown increased T cell mediated cytotoxicity compared to control siRNA cells (\**P* < 0.05, n = 3). **D.** Cytotoxicity in control vs PTEN siRNA transfected DU145 cells. PTEN knockdown significantly decreases cytotoxicity compared to control siRNA cells Chemerin is able to recover overall cytotoxicity in PTEN siRNA cells compared to PBS treated control siRNA cells (\**P* < 0.05 compared to control siRNA + PBS, ΔP < 0.05 compared to PTEN siRNA + PBS, ♦P < 0.05 compared to control siRNA + chemerin, n = 3 independent experiments).

We then investigated the mechanisms of chemerin’s ability to augment T cell-mediated cytotoxicity. Using siRNA knockdown, we again examined the role of CMKLR1 in mediating chemerin’s effects. siRNA significantly abrogated both mRNA and surface protein levels of CMKLR1 in tumor cells (supplemental figure 4). Neither control nor CMKLR1 siRNA affected the low level of cytotoxicity seen using naïve T cells in either PBS or chemerin-treated tumor cells. However, CMKLR1 knockdown completely abrogated chemerin’s effect on activated T cell cytotoxicity, whereas control siRNA had no effect (figure 6B).

### Chemerin augmentation of cytotoxicity is mediated, in part, by PTEN and PD-L1

Given chemerin’s impact on both PTEN and PD-L1 expression, we then explored their roles in the cytotoxicity assay. Control or PD-L1 siRNA were then used to look at the role of PD-L1 in this setting (supplemental figure 7). Again, no effect of siRNA transfections were seen with naïve T cells. Using activated T cells, control siRNA again had no impact on the increase in chemerin-mediated tumor killing, while PD-L1 knockdown significantly increased cytotoxicity in both control and chemerin-treated tumor cells (figure 6C). This is not surprising given the high levels of PD-1 found on the activated T cells (supplemental figure 6), and known effects of blocking PD-L1 in this setting ^56^. Interestingly, levels of cytotoxicity in the PD-L1 knockdown groups were comparable to that in the chemerin/control siRNA groups. Control siRNA + chemerin-treated cells displayed 29% lysis compared to 31% and 33% lysis in the control PBS and chemerin-treated PD-L1 knockdown groups, respectively. While there was a small difference in activated T cell lysis between chemerin-treated control siRNA cells compared to chemerin-treated PD-L1 siRNA DU145 cells, this was not statistically significant. Similarly, there was no significant difference in activated T cell lysis between the PBS vs chemerin-treated PD-L1 siRNA DU145 subsets. We then examined the effects of PTEN knockdown via siRNA transfection (supplemental figure 7). In control-treated cells, knockdown of PTEN resulted in significantly *less* killing by activated T cells, approximately 27% lysis in control siRNA cells compared to 23% lysis following PTEN knockdown. PTEN knockdown in chemerin-treated cells resulted in abrogated T cell killing, to the level of PBS-treated/control siRNA tumor cells at 27% lysis (figure 6D). PTEN knockdown significantly reduced the effect of chemerin treatment – thus, the *difference* in cytotoxicity seen between control and chemerin-treated cells using control siRNA was significantly *greater* than the increase seen using PTEN siRNA. This strongly implicates a mechanistic role for PTEN in chemerin-augmented T cell cytotoxicity. Together, these data support roles for both PTEN and PD-L1 in how chemerin augments sensitivity to T cell cytotoxicity.

### Chemerin treatment is as effective as atezolizumab at augmenting T cell-mediated cytotoxicity

As PD-L1 knockdown does not represent a current clinical scenario in patients, we next evaluated a clinical grade PD-L1 blocking antibody, atezolizumab ^61^. Again, naïve and activated T cells were cultured with control or chemerin-treated DU145 cells. The addition of isotype antibody to the cytotoxicity assay did not impact the beneficial effect of chemerin treatment, as a significant increase in lysis was still seen between control and chemerin-treated DU145 cells (figure 7A). The addition of atezolizumab significantly increased activated T cell-mediated cytotoxicity in the control-treated DU145 cells (figure 7B), consistent with other studies showing the effects of blocking PD-L1 using *in vitro* cytotoxcitiy assays ^62, 63^. There was no statistically significant difference between chemerin-treated DU145 cells with the addition of either isotype control or atezolizumab antibody. Atezolizumab treatment negated the significant difference in lysis between control and chemerin-treated DU145 subsets. Together, these suggest that the addition of atezolizumab is effective in control-treated cells, with basal PD-L1 expression, but added no additional impact in chemerin-treated DU145 cells, where pre-treatment suppressed PD-L1 expression.

**Figure 7.**
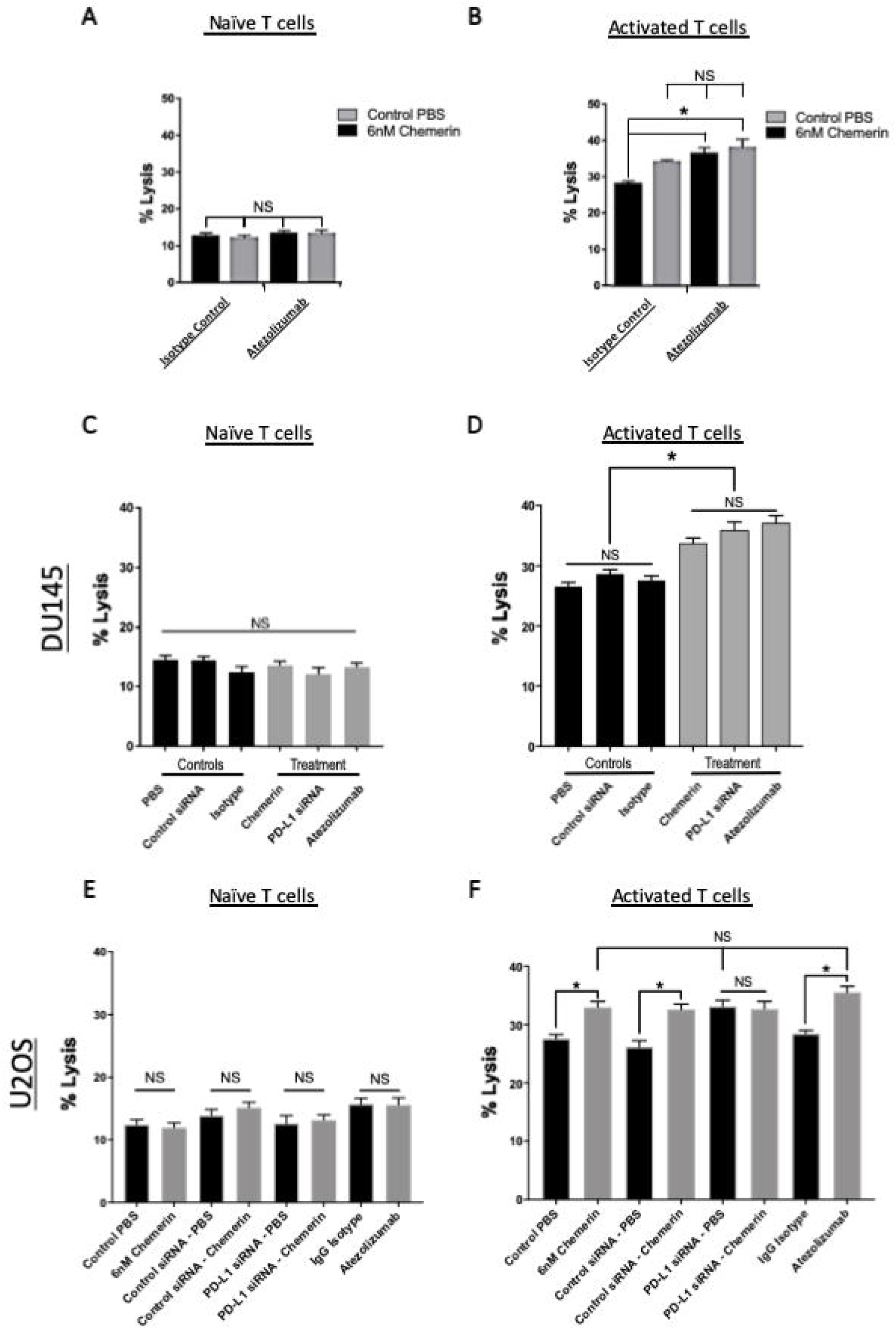
Chemerin treatment increases T cell mediated tumor cytotoxicity comparable to PD-L1 siRNA or Atezolizumab. **A.** T cell mediated cytotoxicity using naive or active T cells against PBS vs chemerin treated DU 145s and IgG isotype. (E:T ratio at 3:1), (\**P* < 0.05, n = 3). **B.** T cell mediated cytotoxicity using naive or active T cells against PBS vs chemerin treated DU 145s and Atezolizumab (anti-PD-L1, 10μg/mL). (E:T ratio at 3:1), (\**P* < 0.05, n = 3). **C.** Cytotoxicity using naive T cells vs. DU 145s treated with the following: PBS, control siRNA, IgG isotype, chemerin, PD-L1 siRNA, or Atezollzumab. (E:T ratio at 3:1), (\**P* < 0.01, n = 3). **D.** Cytotoxicity using 8-10 day active T cells vs. DU145s treated with the following: PBS, control siRNA, IgG isotype, chemerin, PD·L1 siRNA, or Atezolizumab. (E:T ratio at 3:1), (\**P* < 0.01, n = 3). There were no differences in cytotoxicity using naive T cells between each of the respective subsets control vs treatment. For active T cells, Cytotoxicity was significantly increased using chemerin, PD-L1 siRNA, and Atezolizumab compared to their control counter parts: PBS, control siRNA, and lgG lsotype. There was no significant difference in cytotoxicity within the control group and within the treatment group. E. Cytotoxicity using naive T cells vs. U2OSs treated with the following: PBS, 6nM chemerin, control siRNA, PD-L1 siRNA, lgG isotype, or Atezolizumab. (E:T ratio at 3:1), (\**P* < 0 01, n = 3) **F.** Cytotoxicity using 8-10 day activated T cells vs. U2OSs treated with the following: PBS, 6nM chemerin, control siRNA, PD-L1 siRNA, IgG isotype, or Atezolizumab. (E:T ratio at 3:1), (\**P* < 0.05, n = 3).

Subsequently, we compared the T cell cytotoxic effect of chemerin treatment directly to both PD-L1 knockdown as well as Atezolizumab blockade. Using controls and experimental conditions as above, we applied these conditions in parallel, independently repeating this four times with comparable results in both DU145 and U2OS tumor cell lines. No impact was seen in the lower levels of cytotoxicity using naïve T cells (figure 7C, 7E). Using activated T cells, we again found that chemerin treatment significantly increased T cell-mediated cytotoxicity of DU145 cells. Similarly, chemerin exposure to U2OS cells lead to increased T cell-mediated cytotoxicity *in vitro*. Importantly, in comparison to siRNA or Atezolizumab mediated blockade of PD-L1, chemerin treatment was *as effective* at augmenting tumor cell lysis, with no significant differences found between the three conditions in both DU145 and U2OS cells (figure 7D, 7F).

Collectively, these data show that chemerin via CMKLR1 in two different tumor cell lines, DU145 (prostate) and U2OS (osteosarcoma), can induce the upregulation of PTEN and concurrent downregulation of PD-L1 expression in tumor cells. This results in a significant increase in T cell-mediated cytotoxicity, comparable in our assay to the effect of the clinically used Atezolizumab.

## Discussion

Tumor cells have developed various suppressive mechanisms to evade anti-tumor immune responses and regulatory signaling. As both are altered in the TME, further study of the interplay between tumor cell-intrinsic oncogenic signaling and extrinsic anti-tumor immuno-surveillance is necessary to improve current immunotherapies.

The link between PD-L1 (cell-extrinsic immune responses) and PTEN (cell-intrinsic responses) expression has been described, with several examples of PTEN loss or suppression resulting in increased PD-L1 expression in tumors ^45–47, 64^. Other studies suggest PD-L1 expression in prostate, breast, and lung carcinoma may be dependent on PI3k, commonly regulated by PTEN ^64–66^. For example, no association was found between PTEN and PD-L1 expression in two studies of melanoma ^67^ ^35^, supporting the complex and variable regulation of these proteins. However, this association is likely context dependent, as the regulation of PD-L1 expression is controlled by many factors and pathways (figure 8 and reviewed in ^68^). PD-L1 expression correlates with tumor aggressiveness and poor clinical outcomes ^69–75^, as does loss of PTEN ^73, 76–80^, in several datasets, further supporting the clinical impact of alterations in these two key proteins. In prostate, PD-L1 expression has been found to correlate with poorer prognosis and risk of disease recurrence ^73^ ^74^, while PTEN loss has been correlated with both risk of recurrence in localized disease and lethal progression ^80^ ^79^, suggesting a therapeutic strategy to augment PTEN expression may reduce prostate cancer lethality.

**Figure 8.**
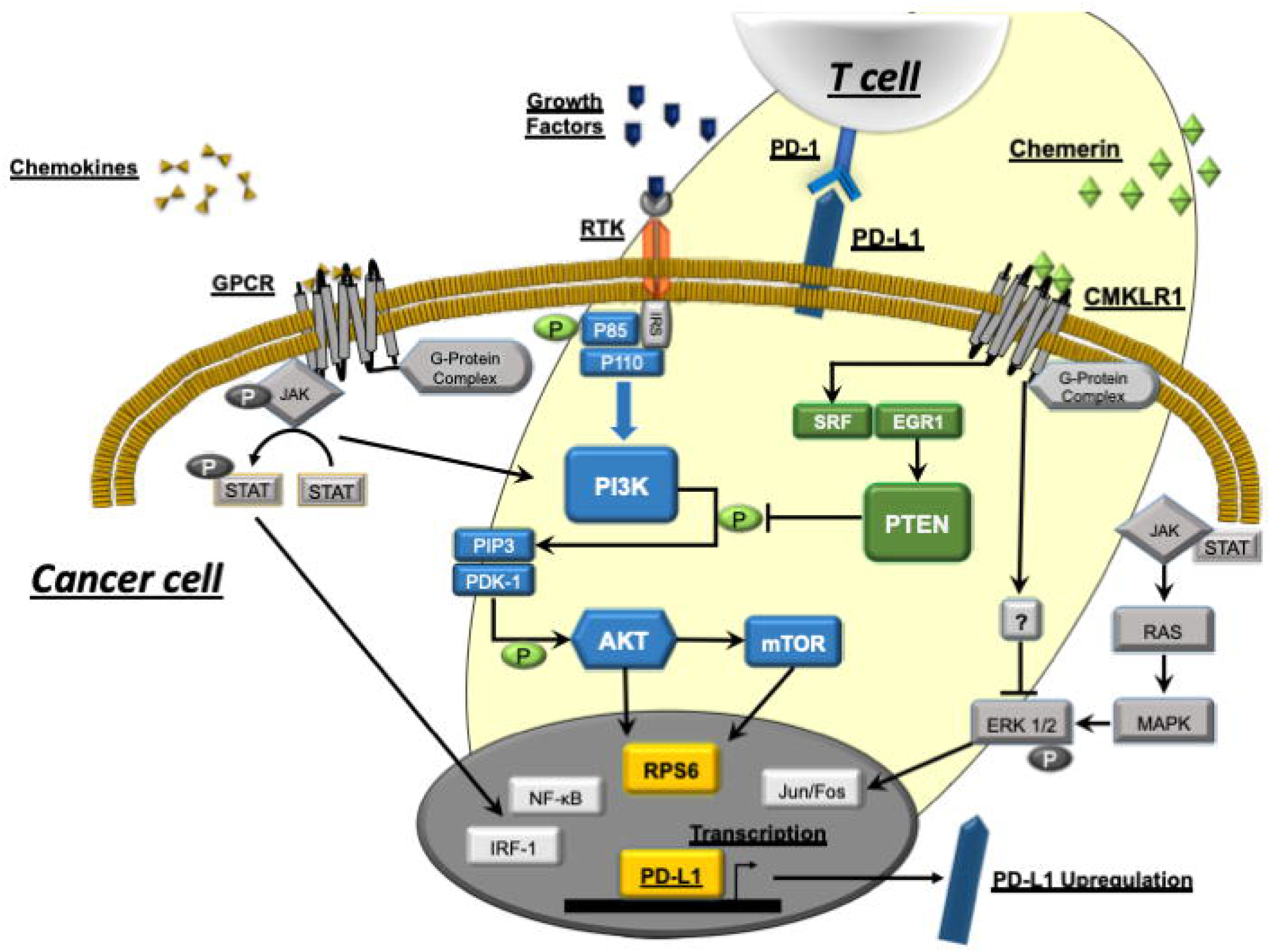
Chemerin/CM KLR1/PTEN/PD-L1 signaling cascade schematic. A current understanding of the Pl3K pathway and how it is regulated by PTEN. Based on our data, we hypothesize chemerin, at east in part, plays a role in modulating PTEN and PD-L1 expression via Chemokine-Like Receptor 1, CMKLR1.

Recent work describes functional consequences of PTEN signaling alterations and its impact on immunoresistance. Toso *et al* used a conditional PTEN null mouse model to study the impact of loss of PTEN within prostate tumors. They found loss of PTEN resulted in a significant increase in several immunosuppressive cytokines, as well as infiltration by CD11b+Gr-1+ granulocytic myeloid-derived suppressor cells (MDSCs) ^34^. More recently, Peng *et al* showed that the PTEN loss led to inhibited T cell-mediated tumor killing and decreased T cell trafficking into the TME. Importantly, they showed that metastatic melanoma patients with PTEN-positive tumors treated with anti-PD1 antibodies had significantly better responses than otherwise matched patients with PTEN-negative tumors. They show that inhibition of PI3Kβ – part of the PI3K/Akt pathway activated with PTEN loss - enhanced the activity of T cell–mediated immunotherapy in mice bearing PTEN-deficient tumors ^35^. Additional evidence recently elucidated the importance of PTEN loss in developed resistance to anti-PD-1 immunotherapy in human sarcoma ^36^, supporting the clinical relevance of this mechanism. Thus, PTEN alterations that impact immunotherapy efficacy are likely one of the important mechanisms to consider in optimization of these therapies.

Our study is the first to show that chemerin, an innate immunocyte chemoattractant, can induce and upregulate expression of functional PTEN in human prostate and sarcoma tumor lines, while concomitantly reducing expression of PD-L1. We show a beneficial functional impact of chemerin treatment in multiple cell lines, with reduced tumor cell migration/invasion and increased T cell-mediated cytotoxicity, on par with the clinically approved anti-PD-L1 antibody atezolizumab.

Importantly, a recent study showed that chemerin could suppress hepatocellular carcinoma (HCC) growth and metastases via the PTEN-Akt signaling axis in a mouse model ^55^. Using human HCC cell lines, Li *et al* showed that chemerin overexpression resulted in upregulation of PTEN and suppression of the PI3K/Akt pathway. As in our studies, they also saw a significant decrease in tumor cell migration/invasion with exposure to chemerin. While not well studied, there is evidence of GPCR signaling leading to increased PTEN phosphatase activity and inhibition of cell migration ^81^, in line with our and Li *et al* ’s data. Similarly, they then used RNA knockdown to show that the effects of chemerin on PTEN and the PI3K/Akt pathway were indeed mediated by CMKLR1. Using a nude mouse model, they showed a significant decrease in *in vivo* HCC growth and metastases. Similar to our findings ^6^, they also found significantly improved survival in patients who had higher chemerin expression within the TME.

Our group was the first to show that augmented chemerin in the TME – often downregulated in human tumors – resulted in recruitment of anti-tumor immune effector cells and significantly reduced tumor growth ^23^. Several other groups have now shown similar findings both in preclinical and clinical scenarios ^6, 55, 82^, supporting the robustness of these data. Our studies elucidate a novel axis between chemerin, CMKLR1, PTEN, and PD-L1 while simultaneously confirming chemerin’s impact on PTEN/PI3k pathway expression ^55^, extending this to additional tumor types (ie prostate, sarcoma). Independent validation of findings across labs and tumor types suggests this axis may be biologically and clinically relevant (figure 8). Further, we identify key transcription factors, SRF and EGR-1, involved in the chemerin-PTEN relationship.

In conclusion, we show a novel role of chemerin in modulating both PTEN and PD-L1 expression and activity via CMKLR1 in human tumor cell lines, resulting in a significant decrease in tumor migration/invasion and increase in T cell-mediated killing. Future studies will explore the impact of modulatiing this axis by chemerin in preclinical models.

## Supporting information

Rennier_Supp Figures

## Acknowledgements

We would like to thank Dr. Brian Van Tine for the generous donation of the sarcoma cell lines used in this publication.

## Contributions

K.R., E.K., G.V., and W.S. performed the experiments and analyzed the data. K.R. and R.P. designed the study and wrote the manuscript. R.P. directed the study and obtained funding. All authors authorized the manuscript.

