## Supplementary material for "Chemerin increases T-cell mediated cytotoxicity of human tumors via modulation of a novel CMKLR1/PTEN/PD-L1 axis": Rennier_Supp Figures

Supplementary Figure 1.

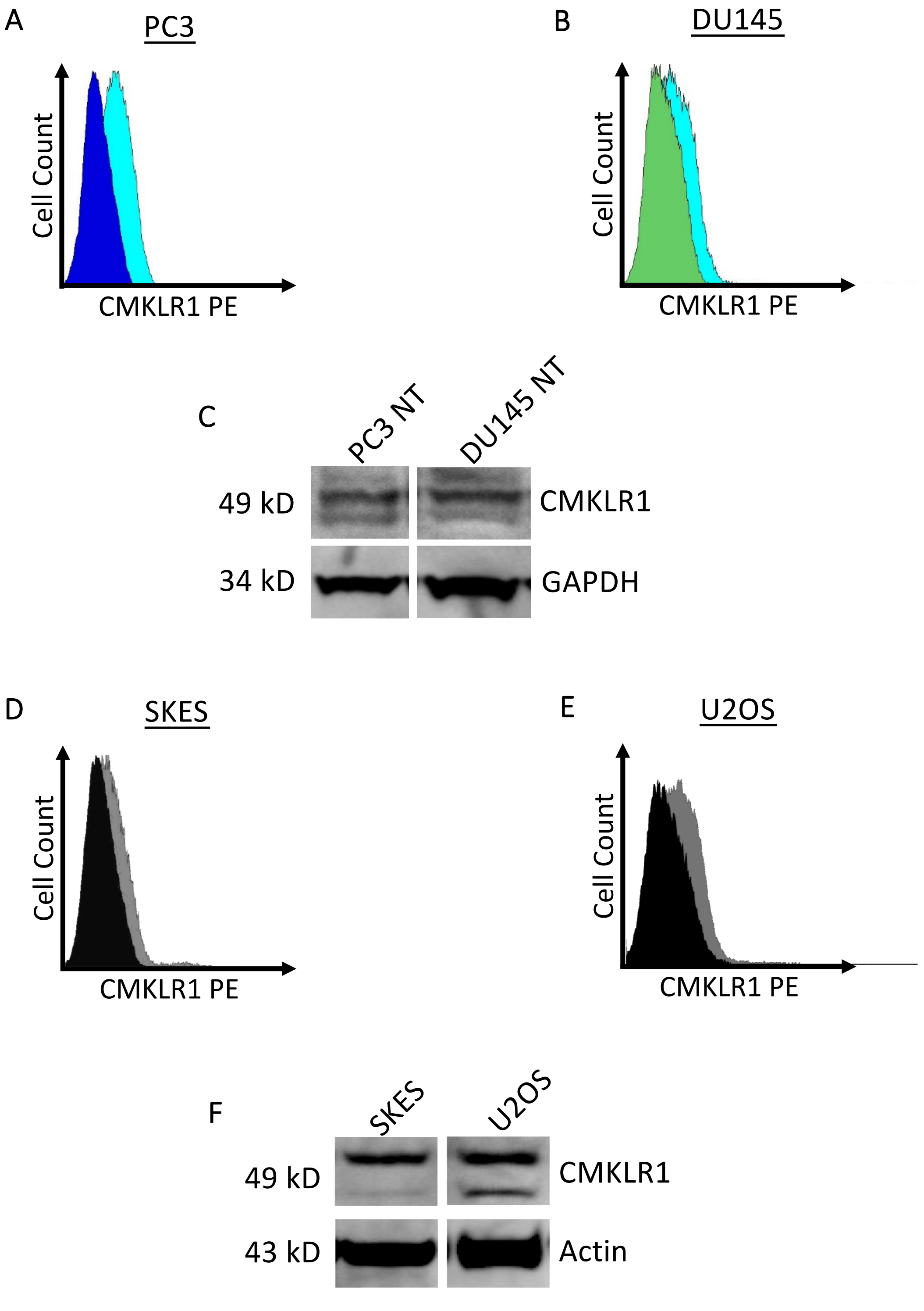


**Supplementary Figure 1. Baseline CMKLR1 Expression in cell lines.** (A) PC3, (B) DU145 CMKLR1 expression measured via FACS (C) Representative Western blot for baseline CMKLR1 expression for PC3 and DU145 cells, normalized to GAPDH loading control. (D) SKES, (E) U2OS cells CMKLR1 expression measured via FACS. (F) Representative Western blot for baseline CMKLR1 expression, normalized to Actin loading control.

Supplementary Figure 2.

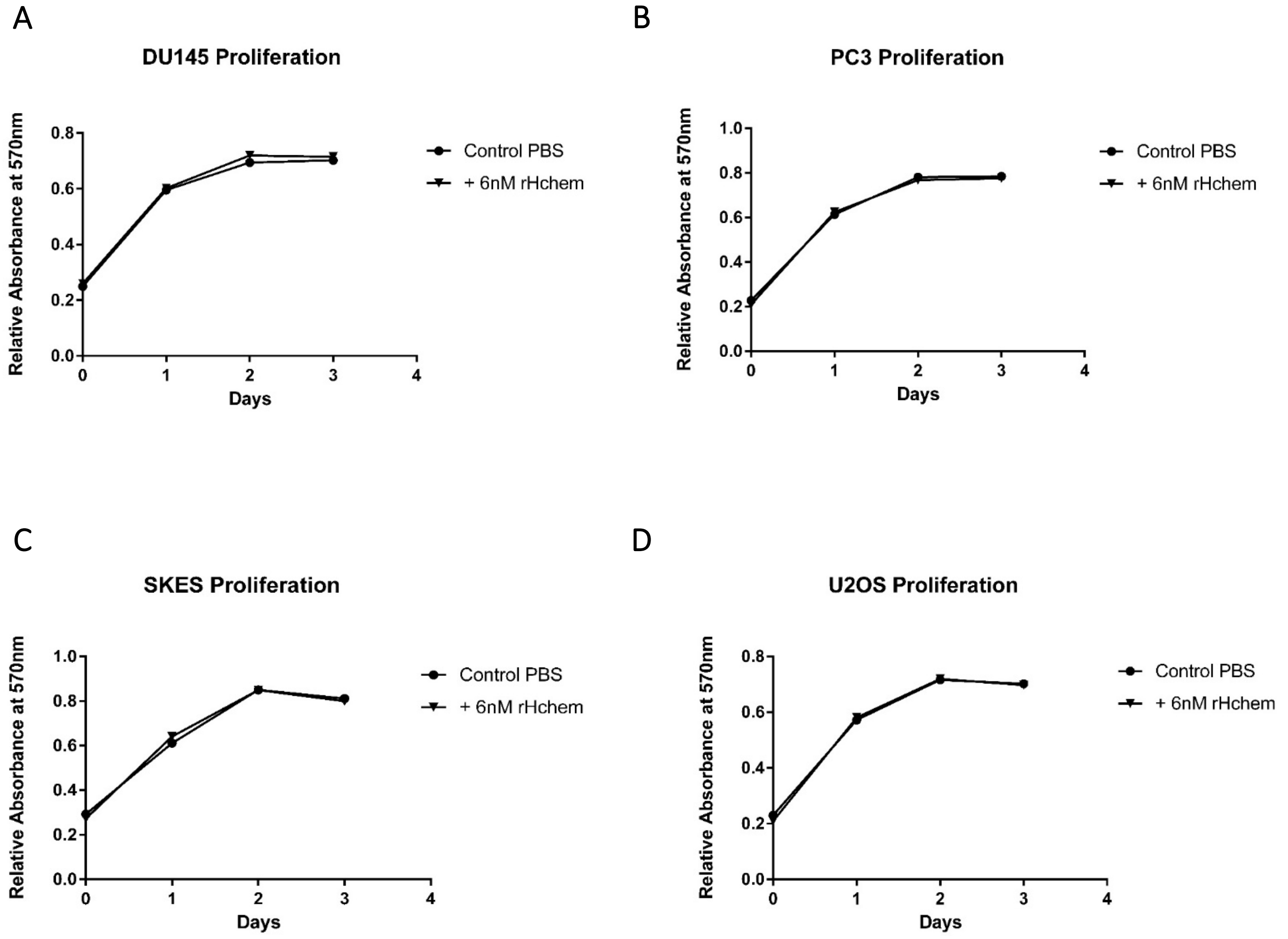


**Supplementary Figure 6. Assessing the effect of chemerin incubation on tumor cell proliferation.** Each cell line was treated with PBS or 6nM chemerin concentration for 72h. Each day, triplicate wells were incubated with Alamar blue, and the absorbance was read, correlating to total number of cells per well (n = 3). Results show that chemerin does not affect overall cell proliferation over a 72h chemerin incubation. *$P < 0.01$, compared to the paired PBS treated cells for each cell line. (A) DU145, (B) PC3, (C) SKES, (D) U2OS.

Supplementary Figure 3.

A

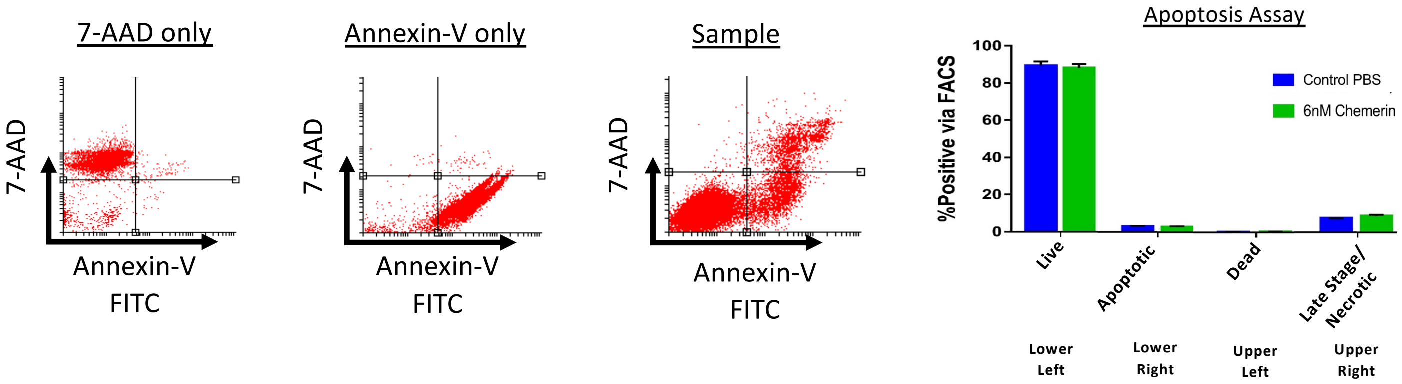


B

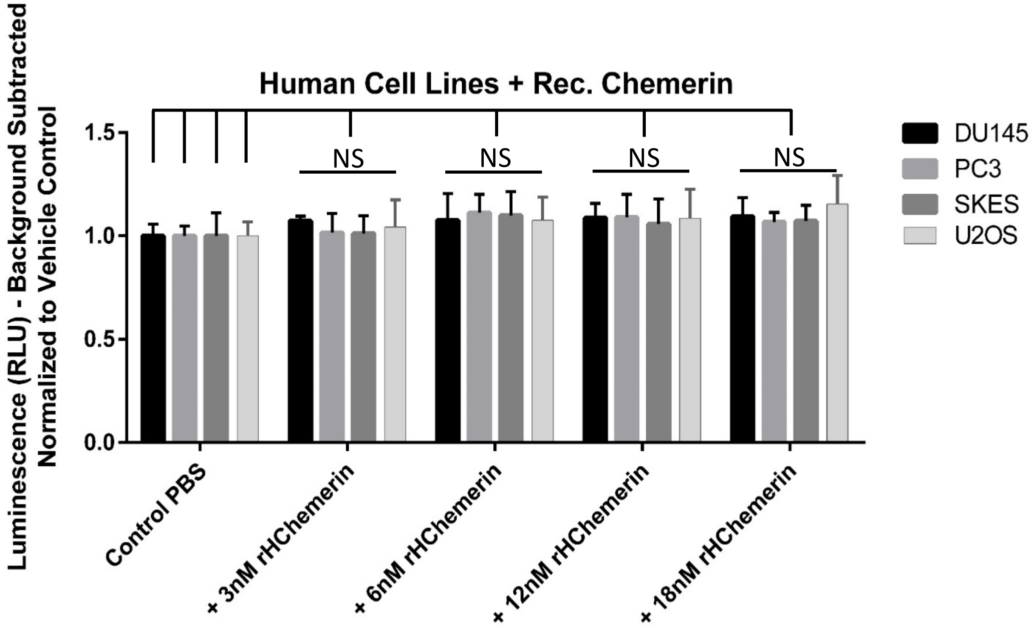


**Supplementary Figure 3. Chemerin incubation has no significant effect on tumor cell apoptosis.** DU145 tumor cells were treated with PBS or 6nM chemerin concentration for 48h. Following initial incubation, cells were continued to be treated for 24h with PBS or chemerin with or without IFN-γ. An Annexin-V FITC antibody was used to stain cells going through apoptosis and 7-AAD stained each cell in the total cell population. Using three independent experiments, triplicate tubes of each cell subset were assessed via FACS (n = 3). **A.** Representative flow cytometry plots showing cells stained with 7-AAD only, Annexin-V FITC only, and a representative sample stained with both 7-AAD vs Annexin-V FITC. Graph of quantitative apoptosis results over 3 independent experiments. There were no significant difference in apoptosis between the PBS vs 6nM chemerin treated cells. (n = 3) **B.** Quantitative graph showing caspase-3/-7 expression in the control PBS vs chemerin treated cells following a 72 hour incubation with vehicle control (PBS) or 6nM chemerin (n = 3). These results show that chemerin does not affect overall cell apoptosis or directly related caspase-3/-7 expression for 72h total incubation in all four cell lines.

Supplementary Figure 4.

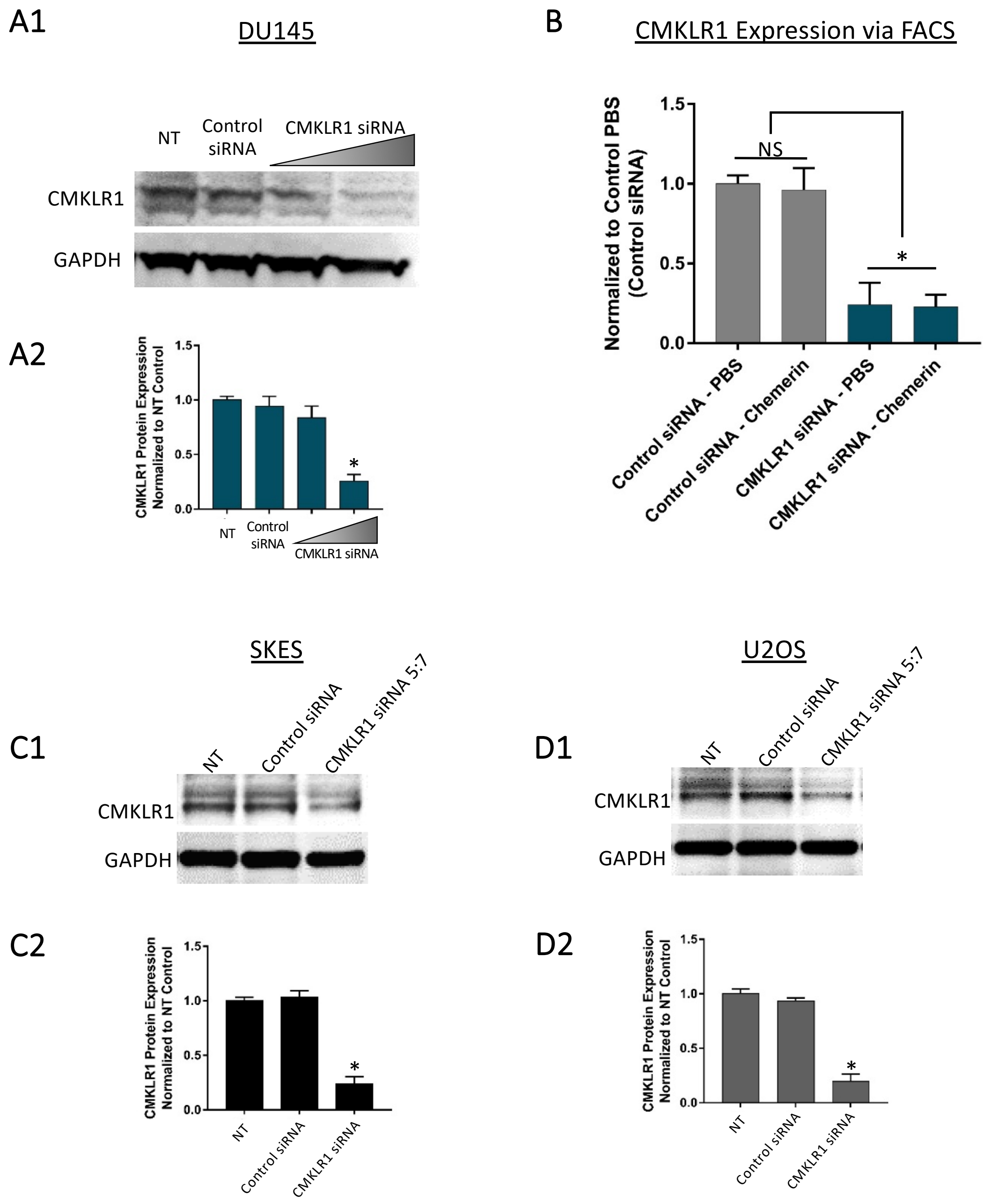


**Supplementary Figure 4. Knockdown of CMKLR1 Expression in Human Cells. A)** Representative Western blot and analysis for CMKLR1 expression after transfection with siRNA, using either 3:9 or 5:7 siRNA to transfection reagent ratio. Loading control bands were probed with anti-GAPDH antibody on the same blot. DU145 (A1, A2). **B)** CMKLR1 receptor expression via FACS in control vs CMKLR1 siRNA (5:7) transfected DU145 cells **C-D)** Western blot analysis for CMKLR1 expression using a 5:7 ratio of siRNA:transfection reagent. SKES (C1, C2), and U2OS (D1, D2) cells were transfected with either control siRNA or siRNA against CMKLR1. Relative band intensity was analyzed based on results from three independent experiments and presented as a ratio to the baseline CMKLR1 expression in non-transfected cells (n = 3). Transfection of CMKLR1 siRNA, but not control siRNA, resulted in a significant decrease in CMKLR1 expression. *$P < 0.01$ compared to non-transfected (NT) cells.

Supplementary Figure 5.

A

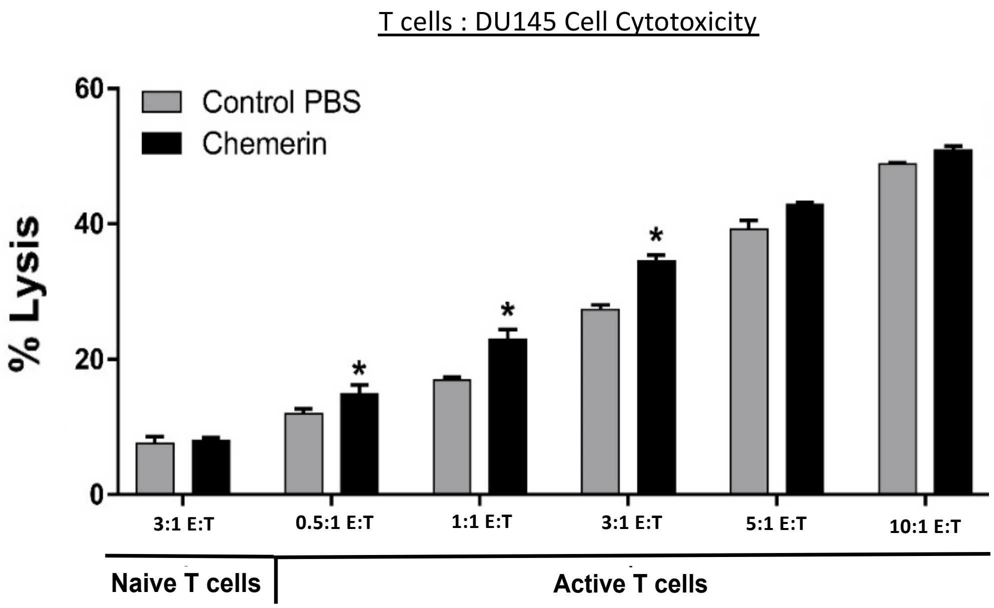


**Supplementary Figure 5. Optimizing effector to target ratio using activated human T cells.** A. Varying effector to target (E:T) ratios showing T cell mediated cytotoxicity in PBS vs 6nM chemerin treated DU145. (**P* < 0.05, using triplicate samples for each experiment and repeated for n = 3 independent experiments).

Supplementary Figure 6.

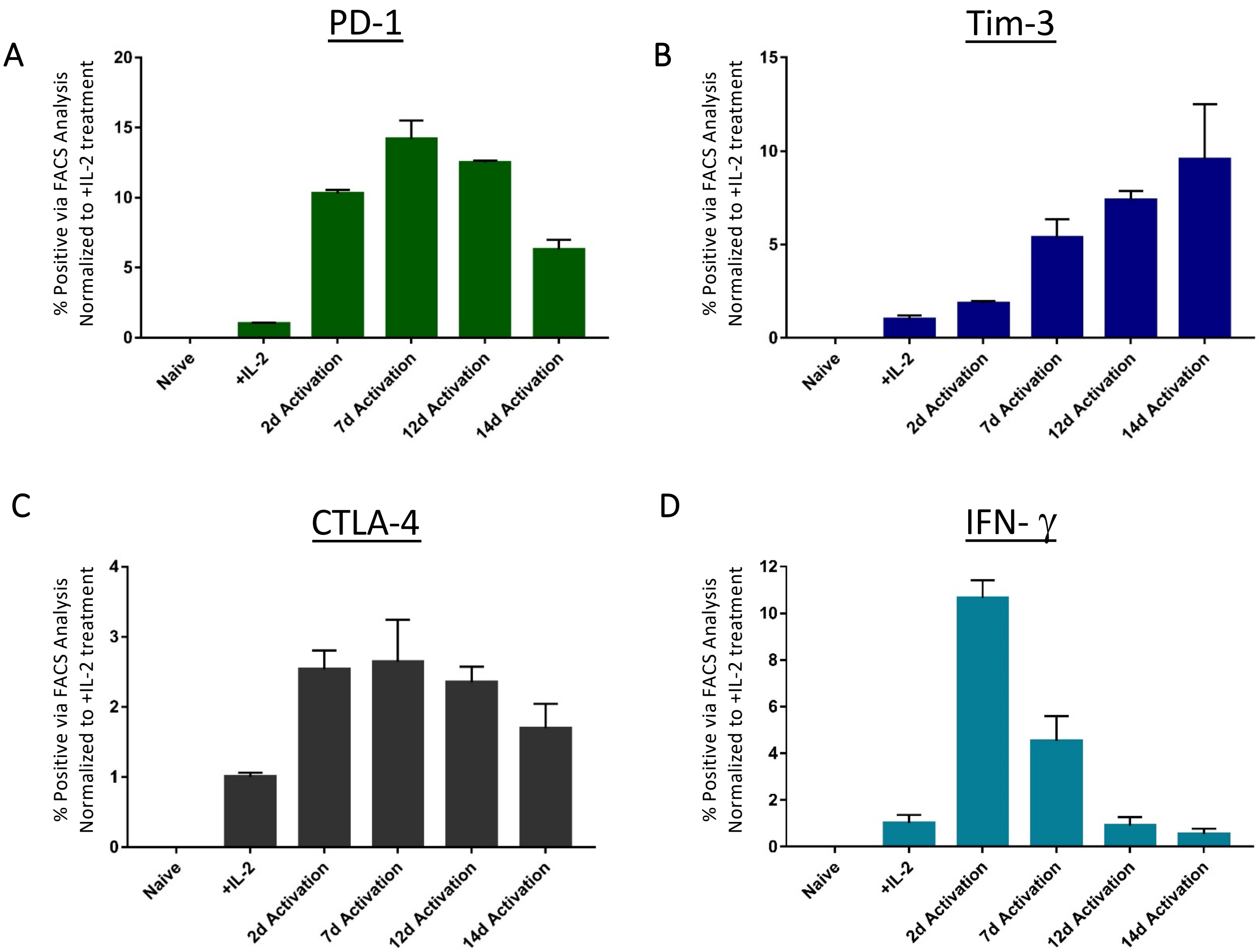


**Supplementary Figure 6. Phenotypic markers on isolated human T cells.** Human T cells were isolated from healthy donor PBMCs. The cells were left untouched (naïve), treated with IL-2 (100U/mL) or exposed to IL-2 + CD2/CD3/CD28 activation tetramers for up to 14 days. Following treatment, cells were assessed for **A.** PD-1, **B.** Tim-3, **C.** CTLA-4, or **D.** IFN-γ expression via FACS (n = 3). Over time, the activated T cells show diminished IFN-γ expression but increased and sustained expression of inhibitory markers: PD-1, Tim-3, and CTLA-4. These results represent a more physiologically relevant, exhausted T cell phenotype necessary to investigate the PD-1/PD-L1 relationship in our experimental setup**.**

Supplementary Figure 7.

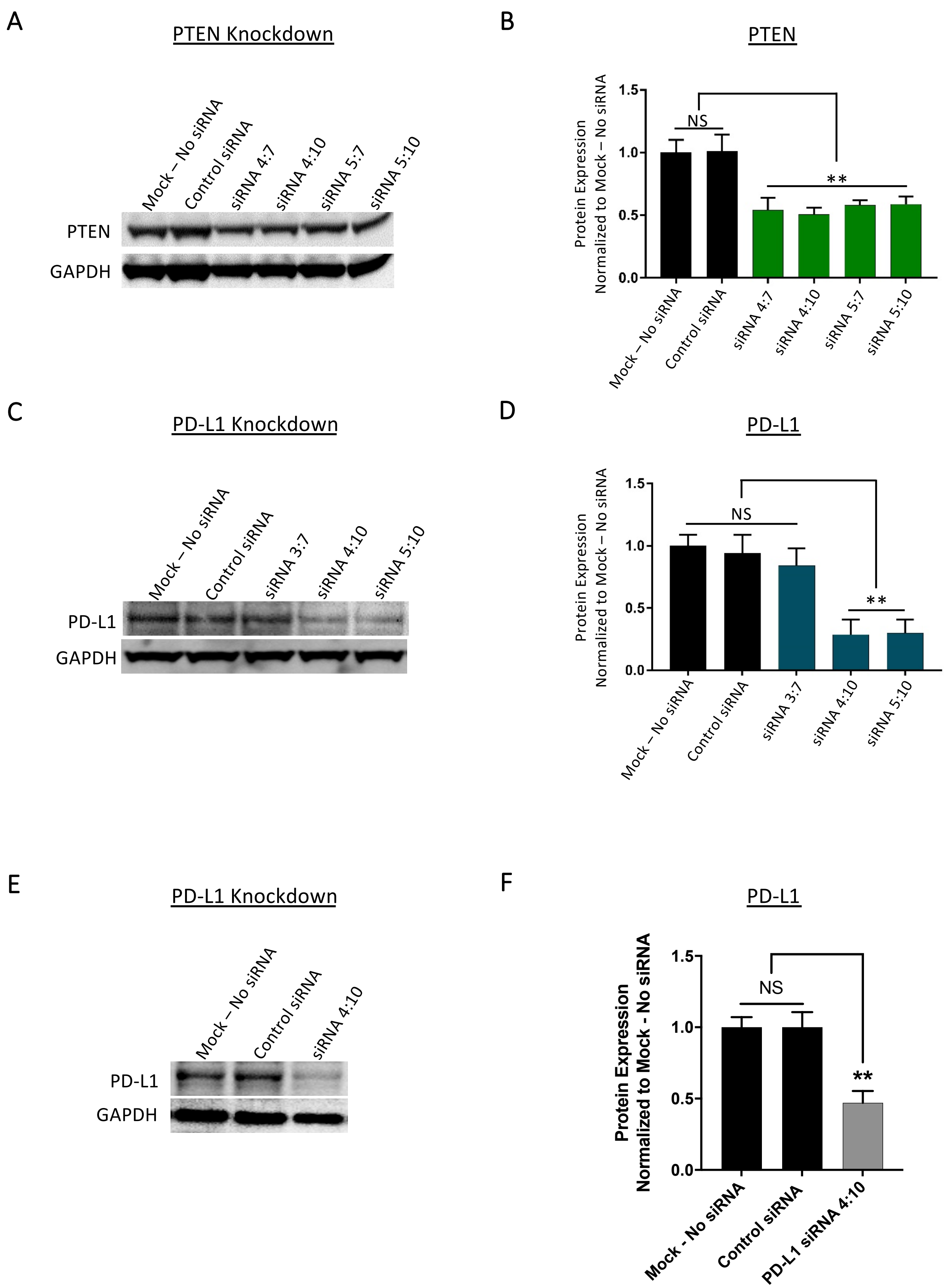

**Supplementary Figure 7. Knockdown of PTEN and PD-L1 Expression in DU145 Cells. A)** Western blot analysis for PTEN expression after transfection with siRNA. Loading control bands are probed with anti-GAPDH antibody on the same blot. DU145 cells were transfected with either control siRNA or siRNA against PTEN. Relative band intensity was quantified and analyzed based on results from three independent experiments, and presented as a ratio to the baseline PTEN expression in mock transfected cells (no siRNA) (n = 3). **B)** Western blot analysis for PTEN expression. Transfection of PTEN siRNA, but not Control siRNA, resulted in a significant decrease in PTEN expression. ***P* < 0.01 compared to Mock – No siRNA cells. **C)** Western blot analysis for PD-L1 expression after transfection with siRNA. Loading control bands are probed with anti-GAPDH antibody on the same blot. DU145 cells were transfected with either no siRNA (Mock), control siRNA or siRNA against PD-L1. Relative band intensity was quantified and analyzed based on results from three independent experiments, and presented as a ratio to the baseline PD-L1 expression in mock transfected cells (n = 3). **B)** Quantified results for PD-L1 siRNA transfection Western blot (n = 3). Transfection of PD-L1 siRNA at the siRNA:transfection reagent ratio of both 4:10 and 5:10, resulted in a significant knockdown in PD-L1 expression, ***P* < 0.05 compared to Mock – No siRNA cells.
